# Comparison of in silico predictions of action potential duration in response to inhibition of I_Kr_ and I_CaL_ with new human ex vivo recordings

**DOI:** 10.1101/2025.02.26.640287

**Authors:** Yann-Stanislas H.M. Barral, Liudmila Polonchuk, Michael Clerx, David J. Gavaghan, Gary R. Mirams, Ken Wang

## Abstract

During drug development, candidate compounds are extensively tested for proarrhythmic risk and in particular risk of Torsade de Pointes (TdP), as indicated by prolongation of the QT interval. Drugs that inhibit the rapid delayed rectifier K^+^ current (I_Kr_) can prolong the action potential duration (APD) and thereby the QT interval, and so are routinely rejected. However, simultaneous inhibition of the L-type Ca^2+^ current (I_CaL_) can mitigate the effect of I_Kr_inhibition, so that including both effects can improve test specificity. Mathematical models of the action potential (AP) can be used to predict the APD prolongation resulting from a given level of I_Kr_ and I_CaL_ inhibition, but for use in safety-testing their predictive capabilities should first be carefully verified. We present the first systematic comparison between experimental drug-induced APD and predictions by AP models. New experimental data were obtained *ex vivo* for APD response to I_Kr_ and/or I_CaL_ inhibition by applying 9 compounds at different concentrations to adult human ventricular trabeculae at physiological temperature. Compounds with similar effects on I_Kr_ and I_CaL_ exhibited less APD prolongation compared to selective I_Kr_ inhibitors. We then integrated *in vitro* IC_50_ patch-clamp data for I_Kr_ and I_CaL_ inhibition by the tested compounds into simulations with AP models. Models were assessed against the *ex vivo* data on their ability to recapitulate drug-induced APD changes observed experimentally. None of the tested AP models reproduced the APD changes observed experimentally across all combinations and degrees of I_Kr_ and/or I_CaL_ inhibition: they matched the data either for selective I_Kr_ inhibitors or for compounds with comparable effects on I_Kr_ and I_CaL_. This work introduces a new benchmarking framework to assess the predictivity of current and future AP models for APD response to I_Kr_ and/or I_CaL_ inhibition. This is an essential primary step towards an *in silico* framework that integrates *in vitro* data for translational clinical cardiac safety.

**Author summary:** Before an investigational drug reaches patients, it is tested *in vitro* to ensure it does not disrupt the heart’s electric activity. This testing often focuses on the drug’s ability to block a specific current called I_Kr_, which, if inhibited, can prolong the heart cells’ action potential duration (APD), which is associated with an increased risk of irregular heartbeats (proarrhythmia). Our study examines how blocking another current, I_CaL_, along with I_Kr_, affects APD. We found that adding I_CaL_ inhibition may mitigate the proarrhythmic effects caused by I_Kr_ inhibition alone. Understanding this balance can improve how we assess the cardiac safety of new drugs, potentially saving promising compounds from being incorrectly discarded. Currently, mathematical models help predict such cardiac responses, but no existing model accurately predicted our findings. Our new data could aid in developing more predictive models in the future. This will contribute to safer drug development and more effective treatments.

## 1 Introduction

The rapid delayed rectifier K^+^ current (I_Kr_) is a major ionic current responsible for the repolarisation of ventricular cardiomyocytes (Sanguinetti *et al*., 1995). Inhibition of I_Kr_ prolongs the action potential (AP) duration (APD) and the QT interval (Redfern *et al*., 2003). Many drugs inhibiting I_Kr_ have been shown to increase the risk of Torsade de Pointes (TdP), a potentially deadly arrhythmia (Redfern *et al*., 2003; Li & Ramos, 2017). Regulatory bodies established guidelines ICH S7B and ICH E14 to prevent the development of new compounds with unacceptable pro-arrhythmic risk (ICH, 2005, 2006). According to ICH S7B, the ability of compounds to inhibit I_Kr_ should be tested *in vitro*. Redfern *et al*. (2003) suggested a “safety margin” such that drugs should have a half-maximal inhibitory concentration (IC_50_) of over 30 times their maximal free therapeutic plasma concentration.

Multiple ion channels affect the TdP risk, notably the inhibition of the L-type Ca^2+^ current (I_CaL_) mitigates the arrhythmogenicity of I_Kr_ inhibitors (Mirams *et al*., 2011). AP models can improve the limited specificity of I_Kr_–centric TdP risk assessment by accounting for simultaneous inhibition of multiple ionic currents (Li *et al*., 2017). The Comprehensive in Vitro Proarrhythmia Assay (CiPA) initiative has encouraged the adoption of biophysically-detailed mathematical AP models as a framework to integrate *in vitro* ion channel data and assess drug-induced TdP risk (Colatsky *et al*., 2016). Yet, AP model predictions of APD changes induced by simultaneous inhibition of I_Kr_ and I_CaL_ have not been validated against human data.

In this study, we measure *ex vivo* the APD at 90% repolarisation (APD_90_) in human adult ventricular trabeculae, with inhibition of I_Kr_ and/or I_CaL_ by 9 compounds (Chlorpromazine, Clozapine, Dofetilide, Fluoxetine, Mesoridazine, Nifedipine, Quinidine, Thioridazine, Verapamil). For each compound, we subsequently use patch clamp data to calculate the percentage of block of I_Kr_ and/or I_CaL_ at the compound concentrations in trabeculae experiments. These numbers are then used as inputs into AP simulations to compare the predictions of 11 *in silico* AP models with the *ex vivo* data.

APD_90_ changes from baseline (ΔAPD_90_) induced by I_Kr_ and I_CaL_ inhibition are linked to QT changes (Johannesen *et al*., 2014). By comparing predictions by existing AP models with the *ex vivo* data, we assess their predictivity in a context relevant to drug development. We thus introduce a benchmarking framework to validate current and future AP models. Therefore, this work can be re-used to help develop predictive models for QT change induced by I_Kr_ and/or I_CaL_ inhibition.

## 2 Methods

### 2.1 Ex vivo action potential acquisition

#### 2.1.1 Sharp electrode recording protocol for data acquisition

Experimental AP data were produced by the AnaBios Corporation, following the methods previously described by Page *et al*. (2016). In brief, trabeculae were extracted from adult human hearts that were not suitable for transplantation, sharp electrodes were impaled in isolated cardiac muscle fibers, their electrophysiological activity was recorded at physiological temperature with vehicle or drugs added. Up to three trabeculae per heart were obtained from the inner endocardial wall of the left (78 trabeculae) and right (4 trabeculae) ventricles. 4 to 15 trabeculae were exposed to the same drug.

In each trabecula, the electrophysiological activity was recorded under baseline conditions, then with three increasing drug concentrations. After the last drug condition, a positive control for APD_90_ prolongation with I_Kr_ inhibition was finally performed with 100 nM Dofetilide addition.

At each drug concentration, each trabecula was paced at 1 Hz for a minimum of 25 min, until voltage recordings were stabilised for at least 2 mins. The stability of APs was assessed qualitatively by the experimenter, based on approximate measurements of the resting membrane potential (RMP), AP amplitude (APA) and APD_90_. After reaching stable recordings, each trabecula was paced at 2 Hz for 3 min then paced again at 1 Hz for 3 min. The experimental protocol for drug administration is shown in Figure 1.

**Figure 1:**
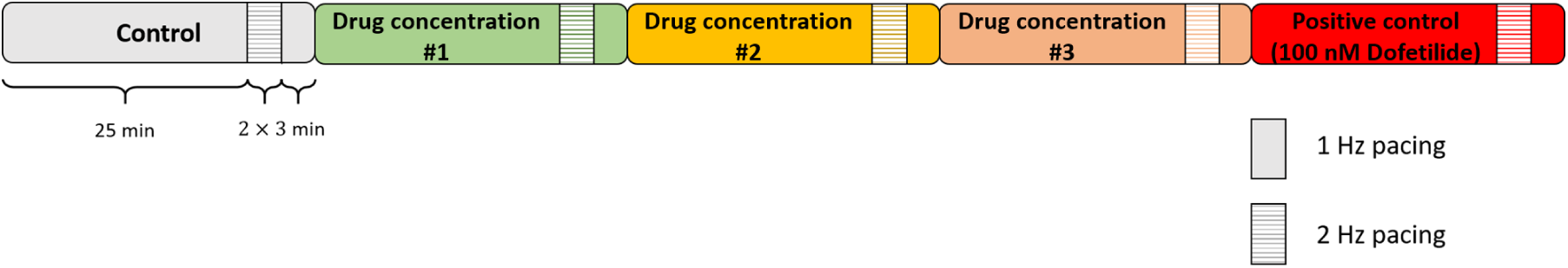
Protocol for sharp electrode recordings of the electrophysiological activity in isolated left-and right-ventricular human trabeculae. After baseline conditions, the response to three conditions with drug was recorded. At the end of the experiments, 100 nM Dofetilide was added as a positive control for APD_90_ prolongation with I_Kr_ block.

For more information on the experimental protocol, see Page *et al*. (2016).

#### 2.1.2 Selected drugs and tested drug concentrations

The tested drugs inhibit I_Kr_ and I_CaL_ with various potencies, so that APD_90_ changes induced by 29 different drug perturbations of I_Kr_ and I_CaL_ could be explored experimentally. The drugs used for this study and their concentrations are reported in Table 1. We call the intended drug concentration (targetted when making up solutions) the *nominal* concentration.

**Table 1:** Drugs tested in *ex vivo* experiments and corresponding nominal concentrations.

| Drug | 1<sup>st</sup> conc<br>(μM) | 2<sup>nd</sup> conc<br>(μM) | 3<sup>rd</sup> conc<br>(μM) | 4<sup>th</sup> conc<br>(μM) | Number of<br>trabeculae |
| --- | --- | --- | --- | --- | --- |
| Chlorpromazine | 0.3 | 1 | 3 |  | 6 |
| Clozapine | 0.3 | 1 | 3 |  | 7 |
| Clozapine | 0.3 | 3 | 30 |  | 4 |
| Dofetilide | 0.001 | 0.01 | 0.1 | 0.2 | 15 |
| Fluoxetine | 0.3 | 1 | 3 |  | 5 |
| Mesoridazine | 0.04 | 0.25 | 10 |  | 6 |
| Nifedipine | 0.003 | 0.03 | 0.3 |  | 4 |
| Quinidine | 0.1 | 1 | 10 |  | 15 |
| Thioridazine | 0.012 | 0.6 | 2 |  | 5 |
| Verapamil | 0.01 | 0.1 | 1 |  | 15 |

Experiments were undertaken in two distinct phases (2014–2016 and 2020–2022). In the second phase (2020–2022), a bioanalysis of the bath solution was performed to measure the drug concentration more precisely at the end of each 25 min period of steady 1 Hz pacing, in case the compound concentration was lowered by absorption by (e.g.) pipettes, tubing or the tissue itself. The sample analysis was performed according to the operating procedure for sample preparation for liquid chromatography–mass spectrometry or mass spectrometry analysis in a bioanalytical laboratory. For data gathered in the first phase (2014–2016), measured concentrations were not available, therefore drug concentrations were assumed to correspond to the nominal concentrations (Table 1).

#### 2.1.3 Ex vivo data post-processing

Voltage was recorded with a time resolution of 0.05 ms and filtered to remove 60 Hz harmonics. The “peak voltage” for calculating percent repolarisation was measured as the upper 95th percentile of voltage (Wang et al., 2015). The resting membrane potential (RMP) was the average voltage over the last 150 ms of the AP. APD_90_ was computed from these reference points and averaged over 30 consecutive APs at the end of steady 1 Hz stimulation. ΔAPD_90_ was defined as the difference from baseline APD_90_. Recordings with 2 Hz pacing were not stabilised after 3 min, so ΔAPD_90_ at 2 Hz was not analysed further.

Sudden changes in resting and peak voltages sometimes occurred, which were attributed to electrode movements: normalized APs showed these did not impact APD_90_, so ΔAPD_90_ was due to drug effects (Barral *et al*., 2022a). Data following voltage discontinuities were discarded if APD_90_ also suddenly altered. Conditions where early after-depolarisations (EADs) were observed (one trabecula with 100 nM Dofetilide) were not analyzed. ΔAPD_90_ was finally averaged over trabeculae exposed to the same drug conditions.

Drug-induced ΔAPD_90_ showed little correlation with baseline APD_90_ — see Supplementary Material D. To align with clinical safety guidelines (ICH, 2006), ΔAPD_90_ was not normalised to baseline and it therefore represents the absolute change from baseline.

### 2.2 Patch clamp measurements of I_Kr_ and I_CaL_ inhibition

Drug effects were modelled as simple pore block, using the Hill equation (Hill, 1910) to characterise it with a half-inhibitory concentration (IC_50_) and a Hill coefficient (*h*). Different voltage-clamp protocols were applied to CHO cells expressing hERG and Ca_V_1.2 channels, and exposed to increasing drug concentrations to measure the drug-induced inhibition of ionic currents.

For I_Kr_ inhibition, two voltage-clamp protocols, denoted “I_Kr_ CiPA” (Kramer *et al*., 2020) and “I_Kr_ Pharm”, were used. Similarly, two voltage-clamp protocols were used to measure the drug-induced I_CaL_ inhibition (“I_CaL_ CiPA” (Li *et al*., 2019) and “I_CaL_ Pharm”). Thereby, two datasets for IC_50_ and *h* for I_Kr_ and I_CaL_ were obtained, denoted the CiPA and Pharm datasets. With these two datasets, the impact of the variability of patch-clamp data (Kramer *et al*., 2020) and of kinetics of drug binding to ion channels (Gomis-Tena *et al*., 2020) can be qualitatively observed. For more details on the generation of the CiPA and Pharm datasets, please refer to Supplementary Material A.

The retrieved IC_50_ and *h* values are reported in Table 2. Dofetilide’s I_CaL_ IC_50_ and Nifedipine’s I_Kr_ IC_50_ were above the highest tested concentration. Therefore, Dofetilide was modelled as a selective I_Kr_ inhibitor and Nifedipine effect as a selective I_CaL_ inhibitor.

**Table 2:** Potency of inhibition of Ca_V_1.2 and hERG channels for drugs tested *ex vivo*. The half-inhibitory concentration (IC_50_) is reported in microMolar (*µ*M), and the Hill coefficient *h* is in brackets. “Pharm” and “CiPA” refer to two different patch-clamp protocols used to characterise I_Kr_ and I_CaL_ inhibition (Supplementary Material A).

| Drug | I<sub>CaL</sub> Pharm | I<sub>CaL</sub> CiPA | I<sub>Kr</sub> Pharm | I<sub>Kr</sub> CiPA |
| --- | --- | --- | --- | --- |
|  | IC<sub>50</sub> (Hill) | IC<sub>50</sub> (Hill) | IC<sub>50</sub> (Hill) | IC<sub>50</sub> (Hill) |
| Chlorpromazine | 2.289 (0.88) | 2.868 (0.93) | 0.359 (0.84) | 0.608 (0.69) |
| Clozapine | 1.676 (0.752) | 4.378 (0.932) | 2.123 (1.05) | 1.978 (0.94) |
| Dofetilide | > 0.2 | > 0.2 | 0.029 (1.10) | 0.033 (1.17) |
| Fluoxetine | 0.994 (0.94) | 0.857 (0.90) | 0.712 (1.26) | 0.772 (0.75) |
| Mesoridazine | 4.056 (0.76) | 3.962 (0.83) | 0.503 (1) | 0.565 (0.74) |
| Nifedipine | 0.105 (0.85) | 0.144 (0.72) | > 8 | > 8 |
| Quinidine | 20.849 (0.63) | 6.68 (1) | 0.966 (1.01) | 0.820 (1.43) |
| Thioridazine | 0.497 (1.07) | 0.637 (1.26) | 0.171 (1) | 0.154 (1.05) |
| Verapamil | 1.381 (0.72) | 0.310 (1) | 0.273 (0.98) | 0.570 (1.67) |

### 2.3 Simulation of APD_90_ with *in silico* action potential models

#### 2.3.1 Selected Models

We selected six main models representative of recent efforts to model the human ventricular AP: Ten Tusscher *et al*. (2004) (TNNP), Ten Tusscher & Panfilov (2006) (TP), Grandi *et al*. (2010) (GPB), O’Hara *et al*. (2011) (ORd), Tomek *et al*. (2020) (ToR-ORd), and Bartolucci *et al*. (2020) (BPS). Since their release, five new parameterisations and variants of these models have been published. Dutta *et al*. (2017) replaced the I_Kr_ component of the ORd model with a 6-state Markov model (Li *et al*., 2017) and rescaled the maximal conductances of five ionic currents (I_Kr_, I_CaL_, I_K1_, I_Ks_, I_NaL_). The GPB, TP and ORd models were rescaled by Mann *et al*. (2016) to capture the effects of I_Kr_ and I_Ks_ inhibition and to reproduce APD_90_ features observed in long QT Syndrome (LQTS) populations. Mann *et al*. (2016) added a late sodium component to their versions of the GPB and TP models, based on the ORd model. Krogh-Madsen *et al*. (2017) proposed a version of the ORd model with rescaled maximal conductance parameters for six ionic currents (I_Kr_, I_CaL_, I_Ks_, I_NaCa_, I_NaK_, I_NaL_) to capture populations with long QT syndrome, which was also included in the present study. All these variant models were included in the present study and are summarised in Table 3.

**Table 3:**
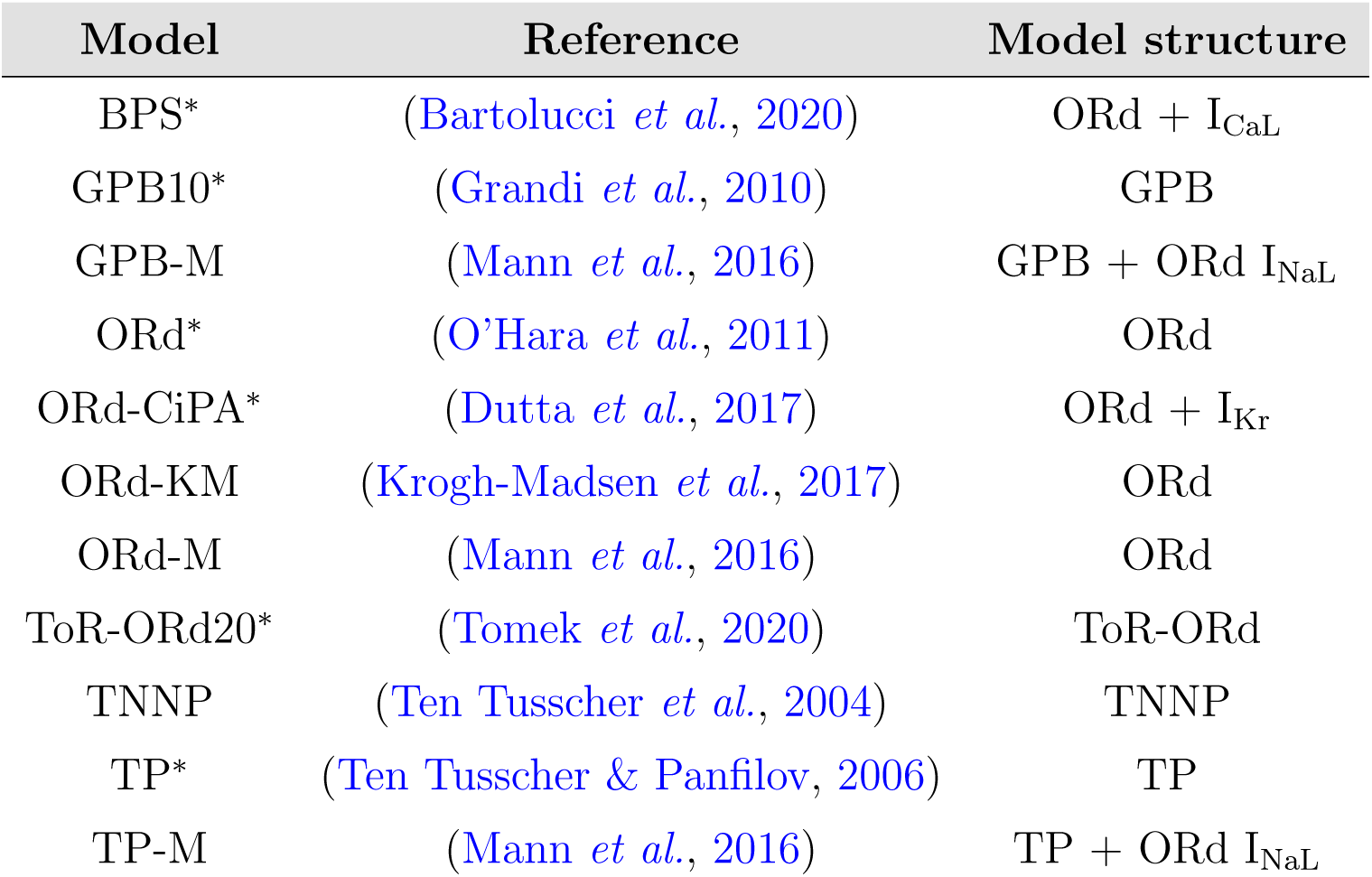
Selected AP models. The ^∗^ symbol indicates when the endocardial version of the model was selected among different versions developed by the authors of the model.

To maximise consistency with the trabeculae measurements, the endocardial variant of the AP models was used.

#### 2.3.2 Action potential simulations

Published CellML models were obtained from the Physiome Repository (Yu *et al*., 2011). Stimulus current width, amplitude, and responsible ions (for instance K^+^(Barral *et al*., 2022b)) were not changed. 1 Hz steady pacing was applied in line with the *ex vivo* experiments. We simulated 1500 s to reach a steady-state response to 1 Hz pacing. In all models, the convergence to steady-state was achieved with 1500 pre-paces (results not shown). The 1501^st^ AP was then recorded with time resolution of 0.05 ms, matching the resolution of the *ex vivo* data, and allowing for precise estimation of APD_90_.

When computing 2-D maps of ΔAPD_90_ as a function of I_Kr_ and I_CaL_ inhibition (Section 3.2), default initial internal and external concentrations from the CellML files were used. When comparing quantitative model predictions with trabeculae recordings (Section 3.3), external concentrations of K^+^, Na^+^, and Ca^2+^ were set to 4 mM, 148.35 mM, and 1.8 mM respectively, matching concentrations used experimentally (Page *et al*., 2016). The steady-state APs simulated with the included models are shown in Figure 2 for visual comparison. Note that the TNNP model does not predict a physiological AP when external ionic concentrations match experimental values. Therefore, predictions with the TNNP model were not quantitatively compared with the *ex vivo* data in Section 3.3.

**Figure 2:**
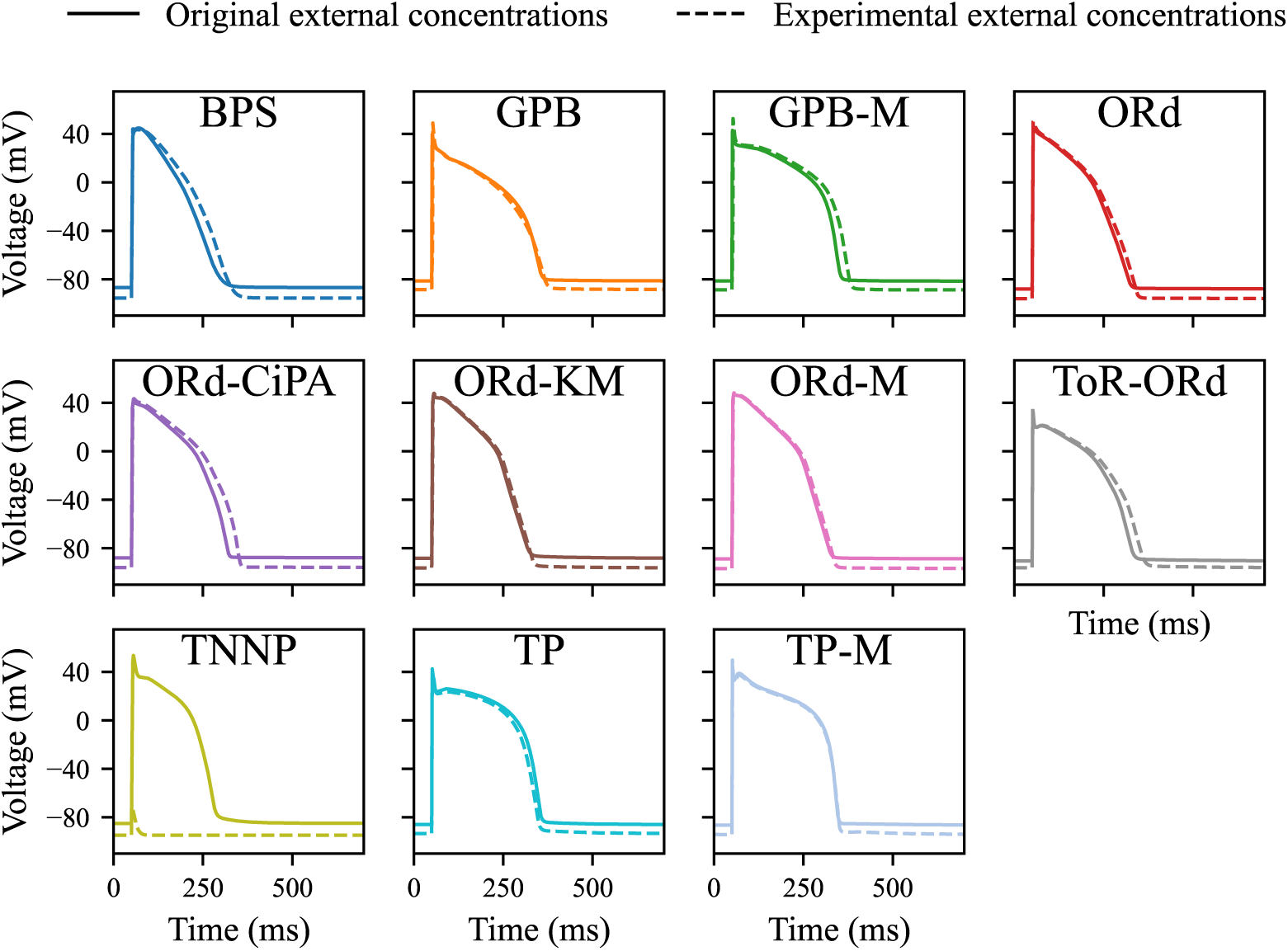
Steady-state 1 Hz AP simulated with the AP models included in this study. External concentrations were set to experimental values (**dashed line**) or left at the values in the original CellML model (**solid line**).

*In vitro* data for inhibition of I_Kr_ and I_CaL_ were integrated into model predictions by applying a rescaling factor, computed with the Hill equation (Hill, 1910; Mirams, 2023), to each affected ionic current:

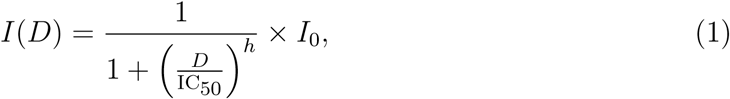

with *I*(*D*) the simulated ionic current, *D* the drug concentration, and *I*_0_ = *I*(0) the ionic current without drug. IC_50_ and *h* were taken from Table 2.

### 2.4 Comparison of model predictive power with experimental action potential data

#### 2.4.1 Qualitative comparison with 2-D maps of ΔAPD_90_ versus current inhibition

With each model, we simulated APs under 101 × 101 = 10, 201 combinations of I_Kr_ and I_CaL_ inhibition conditions, ranging from 0% to 100% inhibition. ΔAPD_90_ was computed for each I_Kr_/I_CaL_ inhibition combination. ΔAPD_90_ was shown using a colour-map that was kept consistent across all the models and which covered the experimental range of drug-induced ΔAPD_90_. Combinations of I_Kr_ and/or I_CaL_ inhibition for which no change in APD_90_ were observed or predicted (∥ΔAPD_90_∥ ⩽ 1 ms) were plotted as white pixels, thus highlighting a “0 ms line”.

The experimental drug-induced ΔAPD_90_ data was reported with similar methods. Additionally, a cubic surface of ΔAPD_90_ as a function of I_Kr_ and I_CaL_ inhibition was fitted through the *ex vivo* data points (details in Supplementary Material B) to facilitate visual comparison of *ex vivo* data with simulations.

Figure 3 shows a schematic visualisation of the methods for 2-D map simulations.

**Figure 3:**
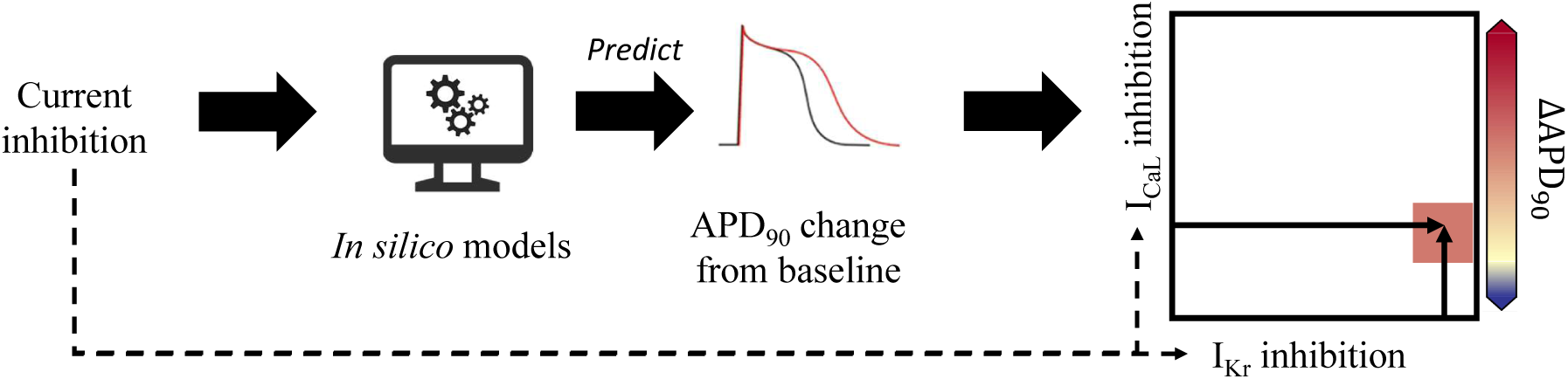
Schematic of methods used to plot ΔAPD_90_ 2-D maps. Simulated ΔAPD_90_ was computed from the *in silico* AP model run for 1500 paces, using I_Kr_ and/or I_CaL_ inhibition as input for the model. The corresponding point was then added to the 2-D map, with ΔAPD_90_ reported with the colour-map.

#### 2.4.2 Metric for the quantification of model predictivity

To quantitatively compare ΔAPD_90_ measurements and predictions, APs were simulated at all nominal drug concentrations, or at actual drug concentrations where available. An error measure, *E*, was then designed to quantify the error in predicted ΔAPD_90_ whilst accounting for experimental variability. *E* was defined as:

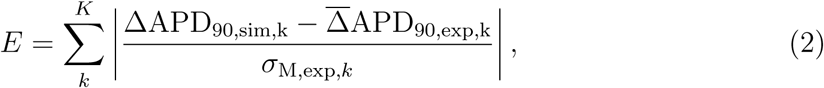

with Δ̅APD_90,exp,k_ and ΔAPD_90,sim,k_ the experimental and simulated ΔAPD_90_ for the drug perturbation *k*, and *σ*_M,exp*,k*_ the standard error of the mean (SEM) experimental ΔAPD_90_ across the trabeculae tested with *k*. Indices *k* span all concentrations of the nine drugs (Table 1).

## 3 Results

### 3.1 Experimental change in APD_90_ from baseline with drug exposure

Experimental APD_90_ measured after 25 min of steady 1 Hz pacing are summarised in Table 4 for the 9 tested compounds.

Chlorpromazine, Clozapine, Fluoxetine, and Mesoridazine induced little or no change in APD_90_ as the effects of I_Kr_ and I_CaL_ inhibition on APD_90_ compensated each other. Verapamil, whilst exhibiting similar effects on I_Kr_ and I_CaL_ (Table 2), substantially shortened APD_90_ (−15 ms to −20 ms on average) with high variability in ΔAPD_90_ (SEM up to 35 ms).

Substantial variability of baseline APD_90_ was observed in trabeculae tested with Mesoridazine, Clozapine, and Nifedipine (SEM of 60 ms, 51 ms, and 55 ms respectively), but this did not lead to particularly high variability in drug induced ΔAPD_90_. The SEM of ΔAPD_90_ was below 7 ms, 9 ms, and 6 ms for Mesoridazine, Clozapine, and Nifedipine, respectively. Fluoxetine-induced ΔAPD_90_ also exhibited low SEM (≤ 7 ms). In contrast, Dofetilide induced the most variable ΔAPD_90_ with up to 33 ms SEM for 200 nM Dofetilide.

The 2-D maps of experimental ΔAPD_90_ are plotted in Figure 4, with I_Kr_ and I_CaL_ inhibition computed with both the CiPA and Pharm datasets (Table 2).

**Figure 4:**
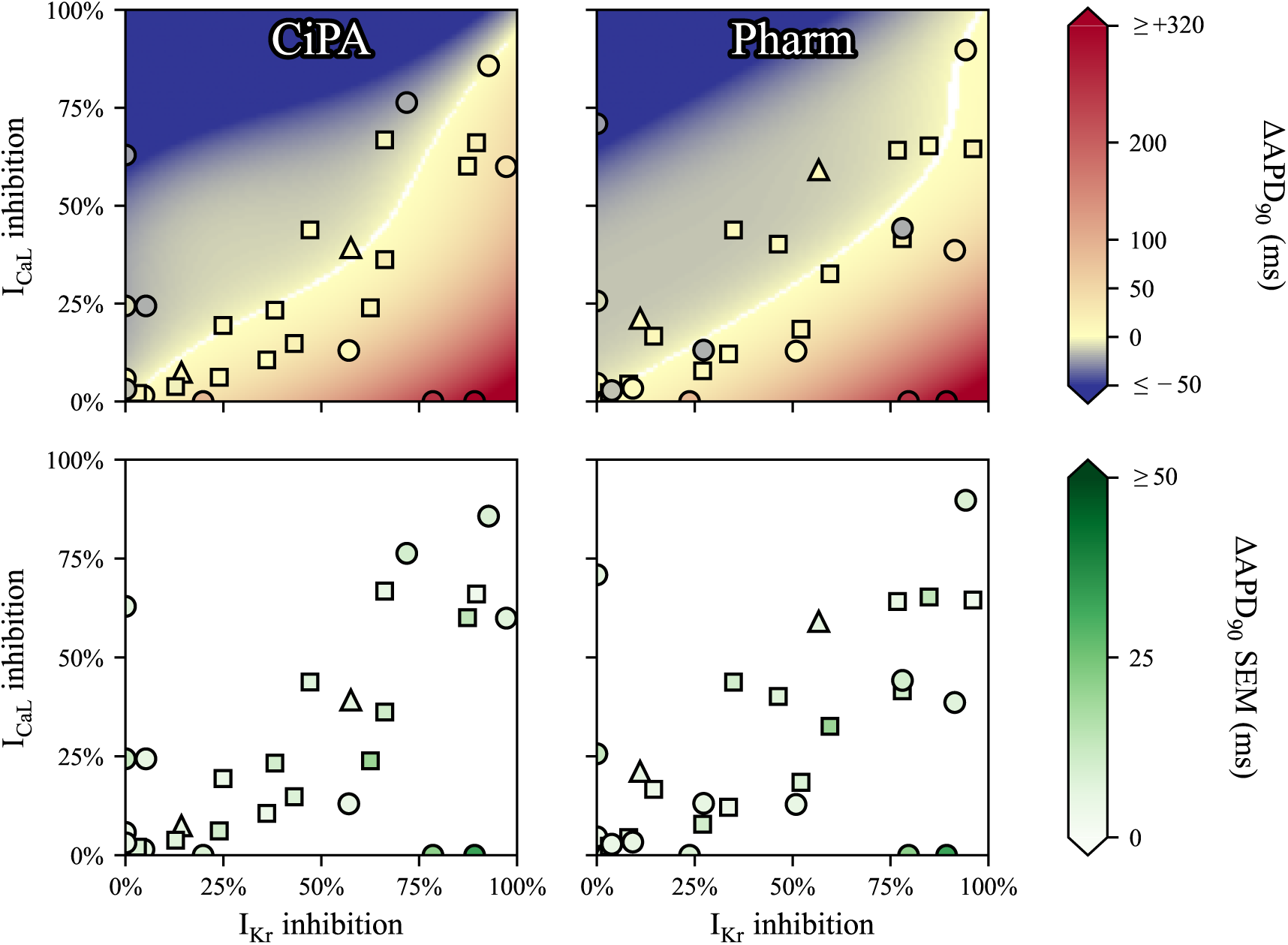
Experimental ΔAPD_90_ measured *ex vivo* under various drug conditions in human ventricular trabeculae, as a function of I_Kr_ and I_CaL_ inhibition and cubic surface approximating the trabeculae data points in the background. I_Kr_ and I_CaL_ inhibition were computed using the Hill equation (Eq. 1), with the CiPA (left) and Pharm (right) datasets reported in Table 2. The bottom panels report the inter-trabeculae variability observed experimentally. When measured drug concentrations were available for all trabeculae tested with the same nominal drug concentration, the data point was plotted as a **square**. When measured drug concentrations were available only for some trabeculae tested with the same nominal drug concentration, the data point was plotted as a **triangle**. When only nominal concentrations were available, the data point was plotted as a **circle**.

With increasing I_CaL_ inhibition, APD_90_ was shortened (shown as darker blue colors). On the other hand, the more I_Kr_ was inhibited, the more APD_90_ was prolonged. I_Kr_ and I_CaL_ inhibition differed from one dataset to another, with the CiPA dataset exhibiting more sensitivity to inhibition of I_CaL_ than the Pharm dataset.

Note that the biggest outlier from the surface, where 1 *µ*M Verapamil induced ΔAPD_90_ = −20 ± 10 ms at 1 *µ*M, with 78% I_Kr_ and 44% I_CaL_ inhibition. For comparison, 3 *µ*M Clozapine induced ΔAPD_90_ = +10 ± 7 ms with 57% I_Kr_ and 59% I_CaL_ inhibition.

### 3.2 2-D maps of ΔAPD_90_ predicted by literature AP models

The 2-D maps of ΔAPD_90_ prediction for all 11 models and variants are shown in Figure 5 with cubic surfaces fitted through experimental data points.

**Figure 5:**
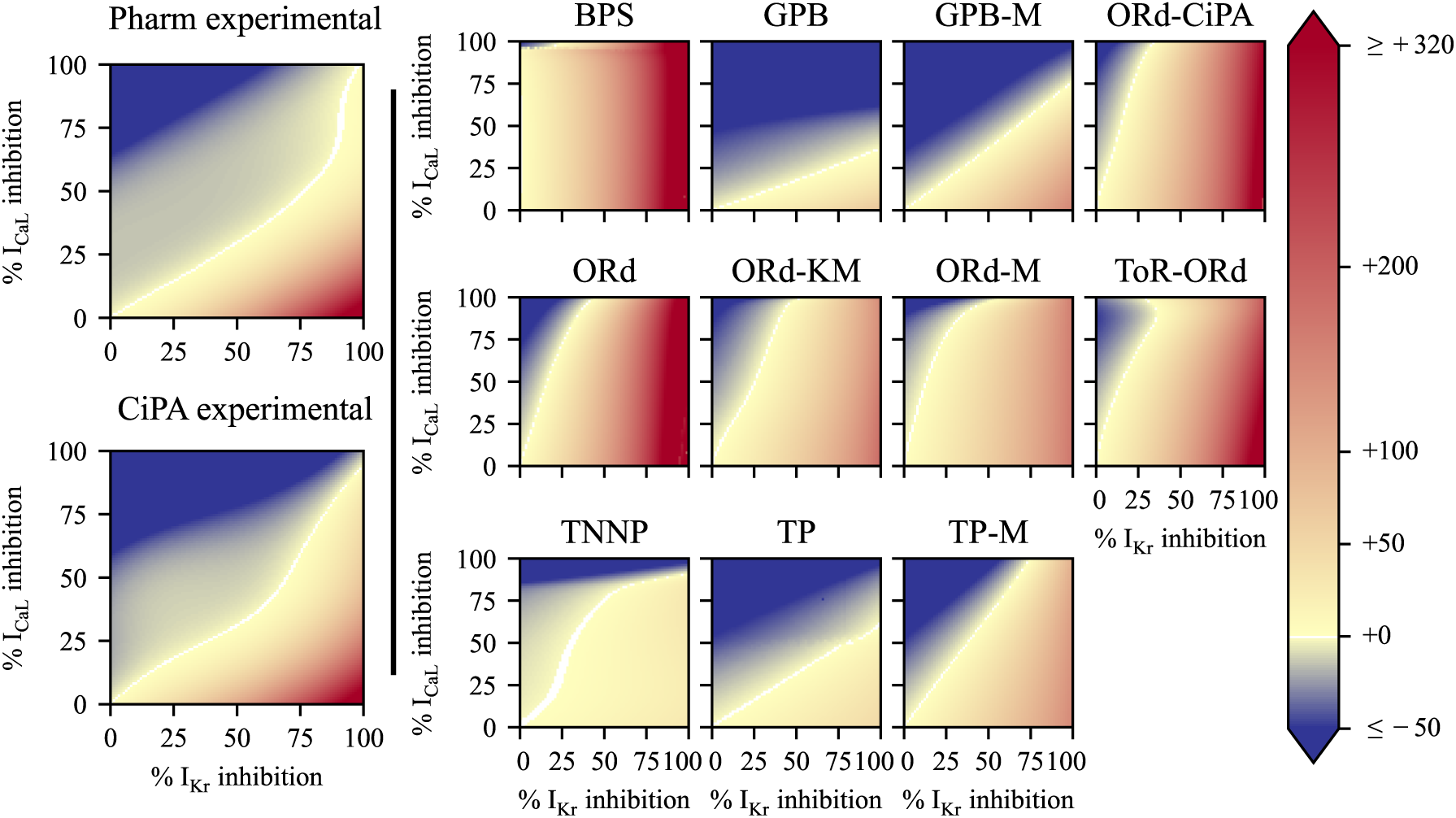
**Left:** Surfaces fitted through experimental data points. **Right:** 2-D maps of predicted APD_90_ change from baseline after I_CaL_ and I_Kr_ inhibition. The colour scale indicates shortening of APD_90_ (i.e., ΔAPD_90_*<* 0 ms) for colours towards dark blue, and APD_90_ prolongation (i.e., ΔAPD_90_*>* 0 ms) for colours towards red. ΔAPD_90_ values below −50 ms and above +320 ms were set to dark blue and red, respectively, for better visualisation. For I_Kr_ and I_CaL_ inhibition leading to −1 ms *<* ΔAPD_90_ *<* +1 ms, the pixel is coloured in white.

A clear distinction was observed between models similar to the ORd model, which were most sensitive to I_Kr_ inhibition, and TP-like models, which were more sensitive to I_CaL_ inhibition. On 2-D maps for the BPS, ORd, ORd-CiPA, ORd-KM, ORd-M, and ToR-ORd models, the 0 ms line was mostly vertical, indicating little mitigation of I_Kr_ inhibition-induced ΔAPD_90_ by I_CaL_ inhibition. These results align with previous observations (Mirams *et al*., 2014).

In the BPS model, nearly no mitigation of I_Kr_ inhibition by I_CaL_ inhibition was observed, and the 0 ms line was vertical. I_CaL_ inhibition even prolonged APD_90_: 5% I_Kr_ and 80% I_CaL_ inhibition yielded ΔAPD_90_ = +9 ms, whilst 5% I_Kr_ and 85% I_CaL_ inhibition yielded ΔAPD_90_ = +11 ms. ΔAPD_90_ predicted by the BPS model was not monotonic. Initially, I_CaL_ inhibition prolonged APD_90_, but with more than 91% I_CaL_ inhibition, APD_90_ decreased drastically.

The ToR-ORd model also exhibited a non-monotonic 2-D map: for 35% I_Kr_ and 90% I_CaL_ inhibition, no change in APD90 was predicted; further I_CaL_ inhibition increased APD_90_. In simulations, the strongly reduced I_CaL_ shrinks the Ca^2+^ concentration in the subspace compartment, reducing the repolarising calcium-activated Cl^−^ current (I_(Ca)Cl_), and therefore prolonging APD_90_.

The TP-like models (TP, TP-M, GPB, and GPB-M) predicted similar 0 ms lines, almost linear with slopes between 0.5 and 1.3. The original TP and GPB models exhibited much lower sensitivities of APD_90_ to selective I_Kr_ inhibition (ΔAPD_90_ ≤ +48 ms and +51 ms respectively), than observed experimentally with 200 nM Dofetilide (ΔAPD_90_ = +318 ± 33 ms). Adjustments by Mann *et al*. (2016) increased their sensitivity to I_Kr_ inhibition: the TP-M and GPB-M models predicted ΔAPD_90_ = +154 ms and +144 ms with 100% I_Kr_ inhibition, respectively.

The TNNP model behaved differently, with its 0 ms line in an “S” shape, and nearly no prolongation of APD_90_, even with 100% I_Kr_ inhibition (ΔAPD_90_ *<* +30 ms).

Visually comparing model predictions with *ex vivo* data, the TP-M and GPB-M models appear closest to the truth. This is quantitatively investigated in the next section.

### 3.3 Comparison of in-silico prediction of ΔAPD_90_ with ex vivo data

Figure 6 presents the quantitative comparison of the *ex vivo* data with ΔAPD_90_ predictions, using external ionic concentrations in simulations that matched the experimental settings. The error in ΔAPD_90_ is shown as a multiple of the experimental SEM in ΔAPD_90_ (*σ*_M_), directly visualising each condition’s contribution to the error measure, *E* (Eq. 2).

**Figure 6:**
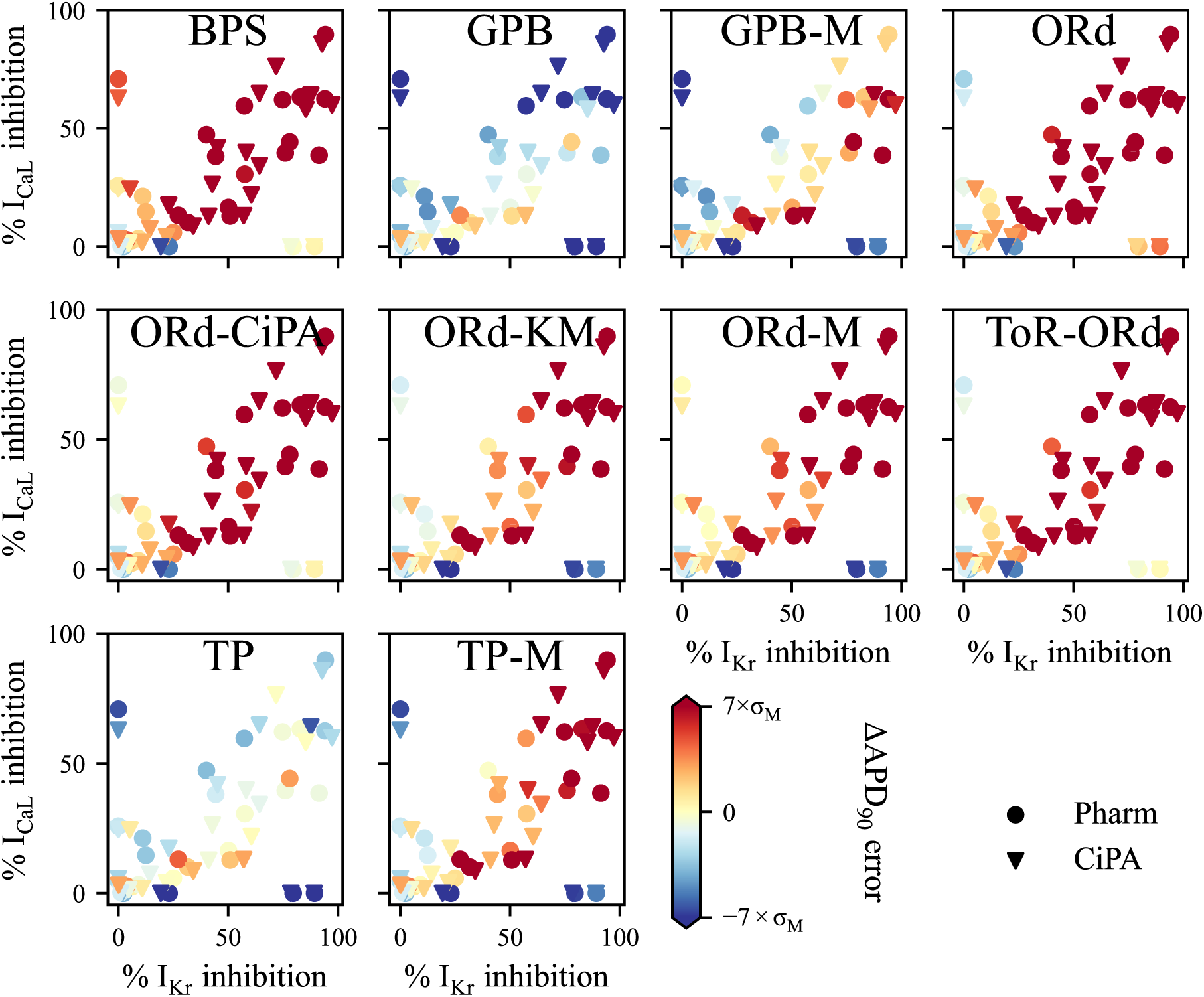
Comparison of *in-silico* prediction of ΔAPD_90_ response to I_Kr_ and/or I_CaL_ inhibition with *ex vivo* data. CiPA (**triangle**) and Pharm (**circle**) datasets for IC_50_ values used to compute drug perturbation are available in Table 2. *σ*_M_ denotes here the experimental standard error of the mean ΔAPD_90_ response to each drug perturbation.

The ORd-like models performed similarly in predicting experimental ΔAPD_90_ consistently with their 2-D maps (Figure 5). Predictions for selective I_Kr_ and I_CaL_ inhibitors by the ORd-CiPA and ToR-ORd models were largely correct (light colors). However, ORd-like models overpredicted APD_90_ prolongation induced by simultaneous I_Kr_ and I_CaL_ inhibition (dark red).

The TP-like models underpredicted the ΔAPD_90_ response to selective I_Kr_ inhibition (blue) but provided good predictions for mitigation by I_CaL_ inhibition. The GPB model predicted excessive APD_90_ shortening after more than 50% I_CaL_ inhibition, but its predictions for simultaneous inhibition of I_Kr_ and I_CaL_ were within 3 × *σ*_M_. The GPB-M and TP-M models overpredicted APD_90_ prolongation induced by simultaneous I_Kr_ and I_CaL_ inhibition, depending on the IC_50_ dataset.

Figure 7A compares *E* (Eq. 2) for all 11 models using the CiPA and Pharm datasets.

**Figure 7:**
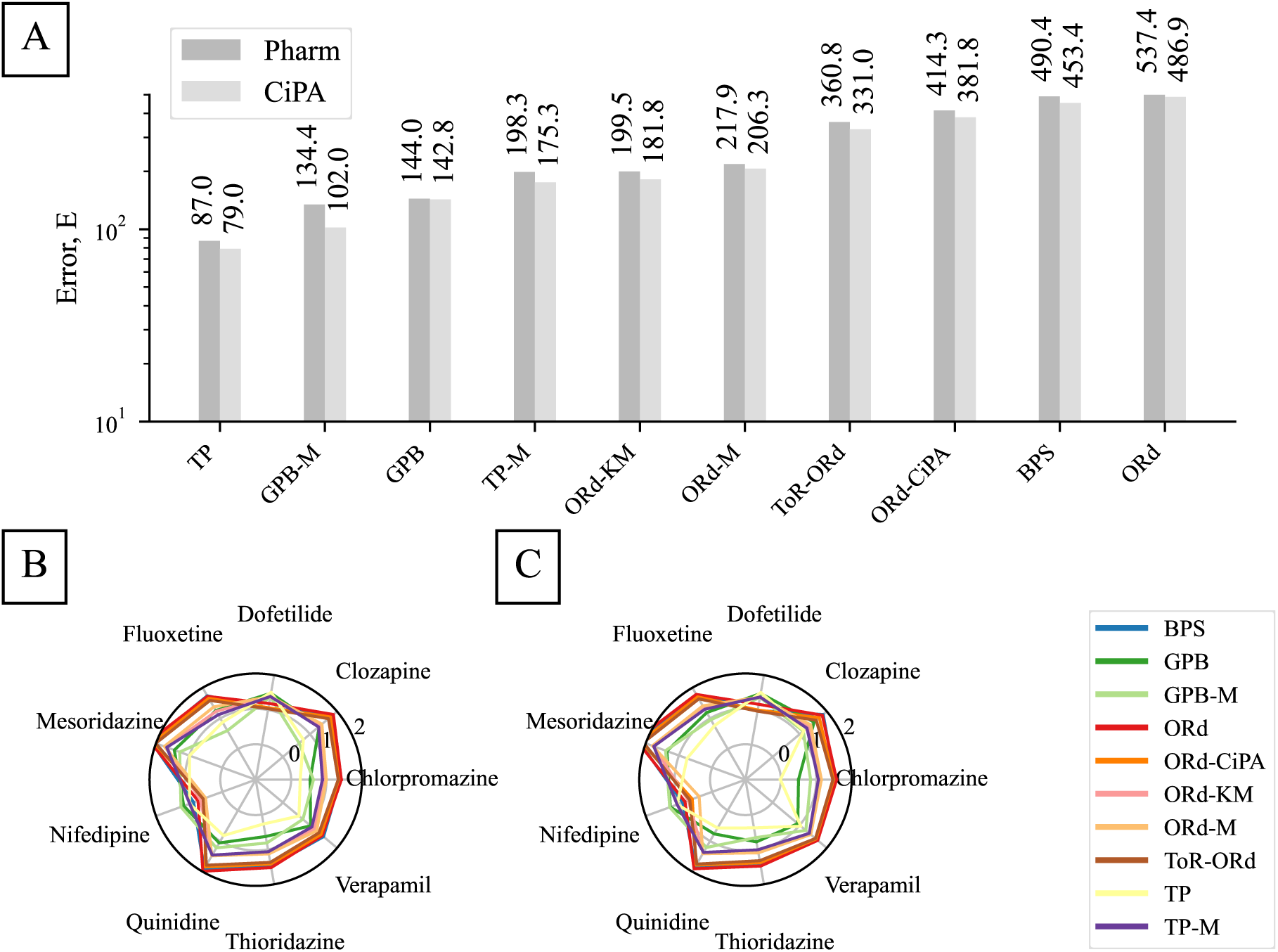
Comparison of the abilities of human ventricular AP models to reproduce the APD_90_ response to I_Kr_ and I_CaL_ inhibition observed *ex vivo*. The lower the error measure (Eq. 2), the more accurate the model predictions. **A:** The error measure was summed over all the drugs used in this study, when using the CiPA and Pharm protocols to compute the reduction of ionic currents by drugs. For each model, two bar plots were plotted, to compare the predictive power of models with the Pharm (left bar) and the CiPA (right bar) datasets. **B and C:** Detail of the error measures associated with each of the drugs using the CiPA and Pharm datasets, respectively, for each model. The log_10_ of the error measure is plotted along the radial-axis.

Figures 7B and 7C detail *E* for each drug.

The TP model yielded the lowest errors *E* = 79.0 and 87.0 using the CiPA and Pharm datasets, respectively. Low errors were found for all drugs with similar effects on I_Kr_ and I_CaL_. The largest *E* for the TP model were for Dofetilide (24.2–31.2) and Nifedipine (8.3–9.8). All TP-like models showed high *E* for Dofetilide and Nifedipine, consistent with Figure 6, where the largest *E* was for selective I_Kr_ or I_CaL_ inhibition.

For the TP and GPB models, *E* computed using the two datasets did not differ significantly. In contrast, models reformulated by Mann *et al*. (2016) and ORd-like models showed stronger dependency on the dataset. A difference of 50.5 was obtained between the CiPA and Pharm datasets with the ORd model: *E* = 486.9 versus *E* = 537.4, respectively. ORd-like models performed similarly, with low errors for Dofetilide and Nifedipine but high errors for other drugs (up to 193.1 for Mesoridazine with the Pharm dataset for the ORd model). These models reproduced the APD_90_ response to selective I_Kr_ or I_CaL_ inhibition well but did not capture the mitigation of I_Kr_ inhibition by I_CaL_ inhibition.

## 4 Discussion

### 4.1 Main findings

The performance of 11 literature AP models was evaluated against new *ex vivo* data from adult human ventricular trabeculae, which measured APD_90_ response to inhibition of I_Kr_ and/or I_CaL_ by 9 different drugs.

The TP-like models exhibited less sensitivity to I_Kr_ inhibition compared to the ORd-like models. The error measure for ΔAPD_90_ prediction, *E* (Eq. 2), was lower with the TP-like models, with the lowest error obtained for the TP model using the Pharm dataset. Their predictions are closer to experimental values for the mitigation of APD_90_ response to I_Kr_ inhibition by I_CaL_ inhibition, but are less accurate for selective I_Kr_ and I_CaL_ inhibitors.

The opposite was observed with ORd-like models. They make accurate predictions of APD_90_ response to selective I_Kr_ inhibition, but do not capture the mitigating effect of I_CaL_ inhibition on I_Kr_ inhibition-induced APD_90_ prolongation.

Mann *et al*. (2016) demonstrated that rescaling maximal conductance parameters of the TP and GPB models and adding a component for I_NaL_ can increase their sensitivity to I_Kr_ inhibition whilst preserving the compensating effects of I_CaL_ and I_Kr_ inhibition on ΔAPD_90_.

In summary, our novel data can be used as a benchmark to assess the predictivity of any future AP model for ΔAPD_90_ response to I_Kr_ and/or I_CaL_ inhibition. Of the currently available models we selected, the TP and GPB-M models showed better overall performance. They therefore appear as promising base models for predicting ΔAPD_90_ response to multi-ion channel inhibitors and subsequent QT changes, upon further development.

### 4.2 Model differences

Several models in this study were validated against previous experimental data for APD_90_ prolongation following I_Kr_ inhibition (O’Hara *et al*., 2011; Grandi *et al*., 2010; Tomek *et al*., 2020; Mann *et al*., 2016; Krogh-Madsen *et al*., 2017; Bartolucci *et al*., 2020). APD_90_ shortening with selective I_CaL_ inhibition was also included in the development of the BPS, ORd, and ToR-ORd models. For instance, the ORd model’s ΔAPD_90_ predictions were validated for APD_90_ response to 70% I_Kr_ inhibition in guinea pig cardiomyocytes (Sanguinetti & Jurkiewicz, 1990) and to 90% I_CaL_ inhibition in rat cardiomyocytes (Walsh *et al*., 2007). whilst the model qualitatively agrees with experimental APD_90_ responses to selective I_Kr_ and I_CaL_ inhibitors, ORd-like models fail to predict ΔAPD_90_ for simultaneous I_Kr_ and I_CaL_ inhibition. Predictions show an I_Kr_-dominated prolongation where *ex vivo* data show mitigation by I_CaL_ inhibition.

This emphasises the need for context-specific model validation (Li *et al*., 2020). For example, the ORd-CiPA model, validated for TdP risk classification (Li *et al*., 2019), tends to overestimate APD response to simultaneous I_Kr_ and I_CaL_ inhibition. Similarly, the BPS model, validated for 100% I_CaL_ inhibition (Bartolucci *et al*., 2020), struggles to predict responses to milder I_CaL_ inhibition.

The TP model, though not validated against current reduction data, showed a low error measure (*E* = 79.0–87.0) but completely failed to reproduce the APD_90_ increase induced by 100 nM Dofetilide (+26 ms predicted vs +256 ± 21 ms experimentally). This is partially due to its significantly higher I_Ks_ maximal conductance (0.392 mS*/µ*F) compared to ORd-like models (from 0.0011 mS/*µ*F in the ToR-ORd model to 0.0196 mS*/µ*F in the ORd-M model), providing greater repolarisation reserve (Roden, 1998).

These findings highlight the importance of thoroughly examining model capabilities. This will help identifying in which context which AP model should (and should not) be used. The Cardiac Electrophysiology Web Lab facilitates this by testing models under various experimental protocols (Cooper *et al*., 2016).

### 4.3 ΔAPD_90_ in the context of proarrhythmic risk assessment

The TdP risk of 28 reference compounds was categorised under the CiPA initiative (Li *et al*., 2017). The Q_net_ metric, simulated with the ORd-CiPA model, predicts TdP risk based on inhibition of major ionic currents (Li *et al*., 2019). The 2-D map for Q_net_ (Figure 16 in Supplementary Materials), computed with similar methods to those for ΔAPD_90_, shows that low TdP risk combinations of I_Kr_ and/or I_CaL_ inhibition qualitatively match the combinations leading to ΔAPD_90_ ≤ 0 ms. This suggests a qualitative agreement between drug-induced ΔAPD_90_ and TdP risk for drugs inhibiting I_Kr_ and I_CaL_.

Quinidine and Verapamil, which both inhibit similarly I_Kr_ and I_CaL_ (Table 2), exerted a mitigated effect on APD_90_. Their ΔAPD_90_ effects align with their effects on the QT_c_ and JT_peak_ interval of the ECG (Johannesen *et al*., 2014). Our new *ex vivo* data suggest that sufficient I_CaL_ inhibition can prevent changes in APD_90_, QT_c_, and JT_peak_ intervals, even at concentrations higher than I_Kr_ IC_50_ — assuming the compound affects cardiomyocytes only through I_Kr_ and I_CaL_ inhibition. This may explain discrepancies between ICH S7B (high risk with I_Kr_ blockade) and ICH E14 (low risk with no QT_c_ change) guidelines, potentially leading to false positives in pre-clinical risk assessments (Vargas *et al*., 2021).

De Ponti (2008) estimated that 60% of new chemical entities inhibit I_Kr_, possibly including useful compounds with I_CaL_ inhibition mitigating the TdP risk. But due to the prevalence of the I_Kr_–centric risk assessment, these compounds are rarely developed. Identifying combinations of I_Kr_ and I_CaL_ inhibition that do not prolong APD_90_ could help develop compounds incorrectly deemed proarrhythmic. However, effects on blood pressure and myocardial contractility due to I_CaL_ inhibition still require attention.

### 4.4 Study limitations

The tested compounds were assumed to primarily affect I_Kr_ and I_CaL_, though literature suggests they may influence other ionic currents (Van Dyke & Scharschmidt, 1987; Zhang & Hancox, 2002; Kramer *et al*., 2013; Crumb Jr *et al*., 2016; Li *et al*., 2019; Barthmes, 2021). Moreover, the drug-binding kinetics may require more complex models than the simple Hill equation used here (Milnes *et al*., 2010; Lei *et al*., 2024). Ionic current response in adult cardiomyocytes may also differ from the response of hERG1a and Ca_V_1.2 expression systems such as those used in the present work (Harchi *et al*., 2018). Refining these modelling assumptions with additional data would improve *in silico* predictions of drug responses.

Most *ex vivo* data were generated from drugs with similar inhibitory effects on I_Kr_ and I_CaL_ (Chlorpromazine, Clozapine, Mesoridazine, Quinidine). These drugs mainly yielded small ΔAPD_90_ responses, highlighting the importance of more detailed risk assessment for multi-channel ‘balanced’ inhibitors. Furthermore, our error measure, *E*, tends to favour models that accurately reproduce minimal APD_90_ prolongation from mixed inhibition. An AP model predicting ΔAPD_90_ = 0 ms for all combinations would score *E* = 59.0, outperforming all models studied here, indicating that *E* alone offers limited model comparison. Combining *E* with 2-D maps of ΔAPD_90_ prediction (Figure 5) and error maps (Figure 6) helps identifying promising models for further refinement.

Concentration measurements were only available for half of the trabeculae, and drug concentrations were generally lower than nominal values, altering the positions of *ex vivo* ΔAPD_90_ data points on our maps. Yet, the fitted cubic surface and model comparisons were not significantly impacted by the use of only nominal concentrations (Supplementary Material C).

## 5 Conclusion

Our new experimental data provide quantitative understanding of the relationship between APD_90_ and acute I_Kr_ and/or I_CaL_ inhibition in adult human ventricular cardiac muscle. Combined with *in vitro* data, they make a valuable benchmark for assessing the performance of *in silico* AP models. Although certain models accurately predict APD_90_ prolongation for selective I_Kr_ inhibitors, they struggle to replicate the mitigating effects of simultaneous I_CaL_ inhibition observed experimentally. The TP and GPB-M models appear to be the most promising starting points for developing more advanced AP models that can account for multi-ion channel inhibition in cardiac safety risk predictions. Of the ORd-like models, the ToR-ORd model exhibits the most promising balance between I_Kr_ and I_CaL_ and its predictivity may be improved upon reparameterisation. Our study emphasises the importance of context-specific validation of AP models: rigorous testing across various ion channel inhibitions and drug concentrations is essential to ensure models can reliably predict cardiac responses. Extended model validation, alongside high-quality experimental data, will ultimately lead to more accurate and reliable *in silico* frameworks for predicting proarrhythmic risk from *in vitro* data. Such frameworks can be vital in improving the specificity of early identification of potential cardiac safety issues in drug development.

## Declaration of Generative AI and AI-assisted technologies in the writing process

During the preparation of this work the authors used ChatGPT (public v3 and internal Roche v4O) in order to enhance general text readability and conciseness, and to debug code. After using this tool, the authors reviewed and edited the content as needed and take full accountability for the content of the publication.

## Data Availability

The data, models, and scripts used to generate the results in this paper are available at the GitHub repository at the address: https://github.com/CardiacModellin g/APD90_ex_vivo_vs_in_silico. Zenodo (permanent archive of Github): https://doi.org/10.5281/zenodo.14791284.

## Author contributions

YSHMB: Conceptualisation, Data curation, Investigation, Visualisation, Writing – Original Draft Preparation. LP: Conceptualisation, Methodology, Supervision, Validation, Project Administration, Writing – Review & editing. MC: Conceptualisation, Methodology, Supervision, Writing – Review & editing. DJG: Conceptualisation, Supervision, Writing – Review & editing. GRM: Conceptualisation, Methodology, Supervision, Visualisation, Writing – Review & editing. KW: Conceptualisation, Methodology, Supervision, Project Administration, Writing – Review & editing.

## Acknowledgements

The authors thank Dr. Abi-Gerges (AnaBios Corporation) for his support with experimental oversight for *ex vivo* data analysis. The authors thank Dr. Bartolucci and Dr. Severi for their help in implementing the BPS model (Bartolucci *et al*., 2020). The authors thank Evgenia Gissinger and Fabian Häusermann (Dr. Polonchuk’s lab) for performing the patch-clamp IC_50_ experiments.

This research was funded in part by the Wellcome Trust [212203/Z/18/Z]. GRM and MC acknowledge support from the Wellcome Trust via a Wellcome Trust Senior Research Fellowship to GRM. For the purpose of open access, the author has applied a CC-BY public copyright licence to any Author Accepted Manuscript version arising from this submission.

## A In vitro measurements of I_Kr_ and I_CaL_ inhibition

For a consistent comparison of the APD_90_ response to drug perturbation predicted by the AP models included in the present benchmark, in some of which the dynamic hERG binding model cannot be implemented, the inhibition of ionic currents was modelled with the Hill equation:

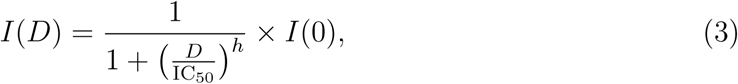

with *I* the current with drug inhibition, *D* the drug concentration, *h* the Hill coefficient, IC_50_ the half inhibitory drug concentration, and *I*(0) the ionic current measured at baseline without any drug exposure.

The AP models considered in this study do not distinguish isoforms of the ion channels, thus the drug inhibition of hERG and Ca_V_1.2 channels was modelled as equal to the drug inhibition on I_Kr_ and I_CaL_, respectively. hERG and Ca_V_1.2 channels are the main ion channels responsible for I_Kr_ and I_CaL_, respectively (Agrawal *et al*., 2022; Sanguinetti *et al*., 1995; Li *et al*., 1996).

The experiments were carried out internally in Roche by Evgenia Gissinger and Fabian Häusermann, from Dr. Liudmila Polonchuk’s lab (Roche Pharmaceutical Research and Early Development, Pharmaceutical Sciences).

### A.1 Cell culture

The CHO crelox hERG cell line was generated and validated at Roche (Guthrie *et al*., 2005). The CHO-hCa_V_1.2/*β*2*/α*2*δ* cell line was purchased from ChanTest (USA, Catalog #CT6004). Vials with cryopreserved cells were thawed at 37°C, washed with the pre-warmed IMDM cell culture medium (Gibco Life Technologies, USA) and re-suspended in the extracellular solution.

For the hERG assay the extracellular solution contained (in mM): NaCl 80; KCl 4; CaCl_2_ 1; MgCl_2_ 1; NMDG 40; HEPES 10; sorbitol 40; glucose 5; pH 7.2–7.4 with NaOH, osmolarity 290-330 mOsm and the internal solution contained (in mM): KCl, 10; KF, 100; NaCl, 10; HEPES, 10; EGTA, 20; pH = 7.0–7.4 with KOH, osmolarity 260-300 mOsm.

For the L-type Ca_V_1.2 assay the extracellular solution contained (in mM): NaCl 80; KCl 4; CaCl_2_ 1.8; MgCl_2_ 1; NMDG 40; HEPES 10; sorbitol 40; glucose 5; pH 7.2–7.4 with NaOH, osmolarity 290-330 mOsm and the internal solution contained (in mM): KCl, 10; KF, 100; NaCl, 5; HEPES, 10; EGTA, 10; Na-ATP, 4; Na-GTP, 0.1; pH = 7.0–7.4 with KOH, osmolarity 260–300 mOsm.

### A.2 Electrophysiology recordings

The recording of the currents was performed using automated patch clamp system SynchroPatch 384 (Nanion Technologies GmbH, Germany) at 35–37°C following the experimental procedure described below. On the day of the experiment, an aliquot of the cell suspension in a 2:1 mixture of the HBSS and external solution was placed in the Cellhotel. The cells were subsequently added into the 384-well sealchip where the currents were recorded in single cells with the patch-voltage-clamp technique in the whole-cell configuration at 35–37°C using the built-in 384 channel amplifier and associated software (PatchControl 384). Currents were low-pass filtered using the analog 3 kHz Bessel filter and the digital 3 kHz Lanczos filter and were digitized at 5 kHz. Series resistance was typically 2–9 MΩ and is compensated by 80%. The reported current amplitudes represent the maximal amplitude of a peak current.

### A.3 Voltage-clamp protocols

The voltage-step protocols used to measure the Ca_V_1.2 inhibition by drugs are shown in Figure 8. In the Roche in-house protocol (Pharm dataset), the cells were held at a resting potential of −90 mV, where the baseline current amplitude was recorded. The Ca_V_1.2 channels were then activated with 120 ms-wide steps at 0 mV with a frequency of 0.1 Hz, where the peak current was recorded. Once the recorded peak and baseline currents were stabilised, the amplitude and kinetics of the I_CaV1.2_ were recorded for 3–5 minutes without drug. Then, after each drug addition, the activity of the cells was recorded during 3 minutes.

In the CiPA protocol, cells were held at a resting potential of −80 mV, where the baseline current was recorded. Then a stimulus 40 ms step of voltage at 0 mV was applied to record the peak current, with a 0.1 Hz frequency. Then the voltage was held at +30 mV for 200 ms, followed by a ramp down of voltage to return to −80 mV in 99 ms.

**Figure 8:**
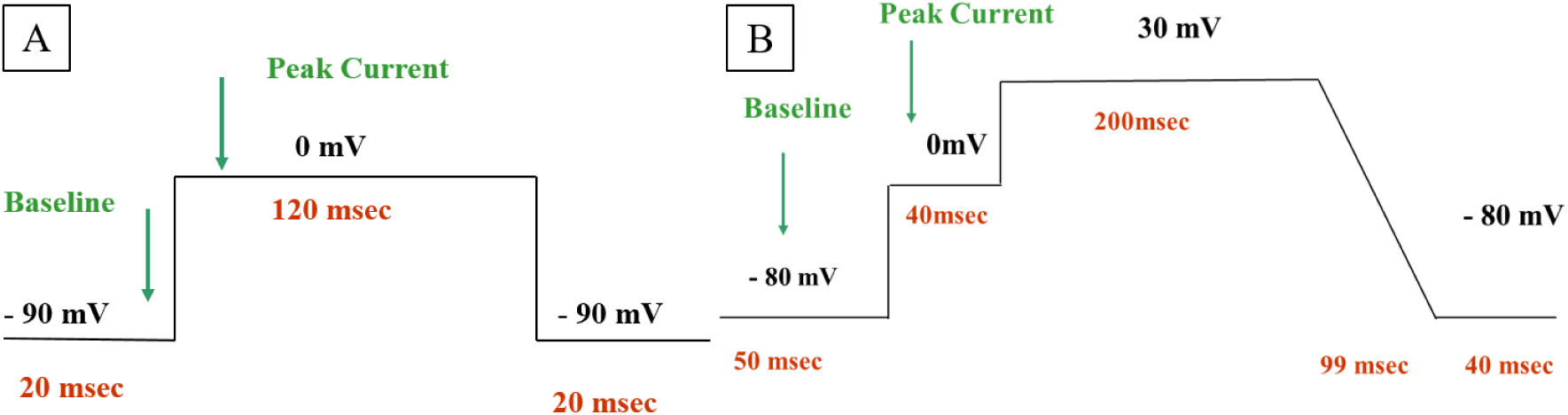
Protocols for recording of the peak I_CaV1.2_ current. **A:** Roche in-house protocol (‘Pharm’ dataset). **B:** CiPA protocol (‘CiPA’ dataset) (Li *et al*., 2019).

The voltage-step protocols used to measure the hERG inhibition by drugs are shown in Figure 9. In the Roche in-house protocol, the resting voltage was set to −80 mV. Then the voltage was clamped to −40 mV for 100 ms, where the baseline current was recorded. The voltage was then brought to +20 mV for 500 ms and finally to −40 mV for 500 ms, where the peak current was recorded. Afterwards, the voltage was set back to the resting potential of −80 mV. The stimulation pattern is repeated with a frequency of 0.1 Hz.

In the CiPA protocol, the cells were held to the resting potential of −80 mV. A +40 mV voltage was then applied to them for 500 ms, followed by a −1.25 mV/ms ramp that brought the voltage down to the resting potential in 96 ms. The peak current was recorded during this ramp down. The pattern was repeated with a frequency of 0.1 Hz.

**Figure 9:**
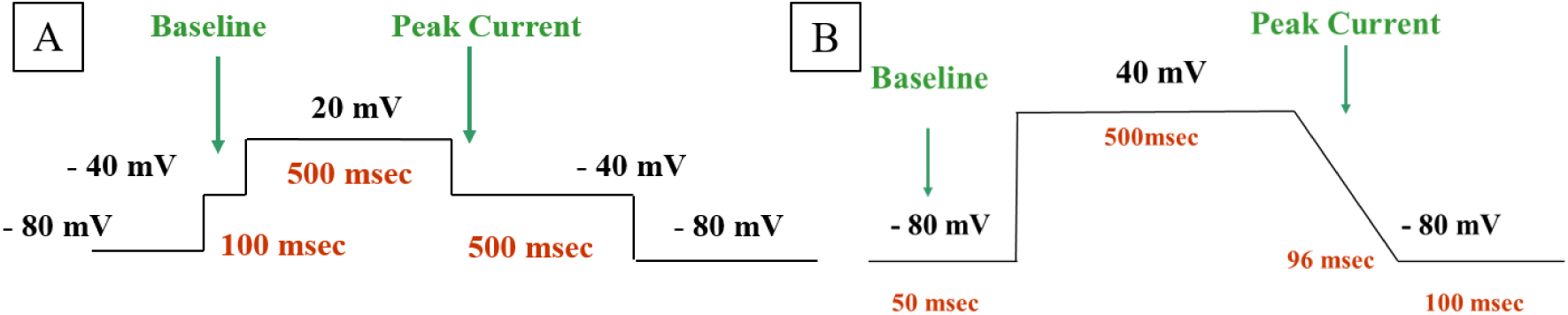
Protocols for recording of the peak I_hERG_ current. **A:** Roche in-house (‘Pharm’ dataset). **B:** CiPA protocol (‘CiPA’ dataset).

### A.4 Literature and new IC50 data

The Tables 5 to 9 below summarise the IC_50_ values that could be found in the literature for hERG and Ca_V_1.2 block potency of Clozapine, Dofetilide, Nifedipine, Quinidine and Verapamil. Overall, there was good agreement between literature values and the values we present in this study. Nifedipine IC_50_ (measured with both protocols) was however higher than literature values for Ca_V_1.2. The 1.5 inter-quartile range statistical test was performed to identify outliers, and no outlier was found.

## B Fitting of the cubic surface through the experimental ΔAPD_90_ data

Cubic surfaces were fitted through the experimental data for drug-induced ΔAPD_90_. The cubic surface was computed following the equation:

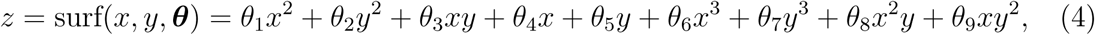

with *z* the approximated ΔAPD_90_, *x* and *y* the percentage of inhibition of I_Kr_ and I_CaL_, respectively, and ***θ*** the parameters describing the cubic surface. The location of each data point for each tested trabecula (*x* and *y*) was computed using the IC50 data (Table 2) and the measured drug concentration in the bath solution if available. The nominal drug concentration was used otherwise.

The cubic surface was fitted through the experimental ΔAPD_90_ data points, by minimising the cost function:

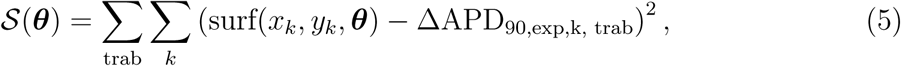

with ΔAPD_90, exp, k, trab_ the experimental ΔAPD_90_ for the drug perturbation *k* averaged over 30 consecutive APs in the trabecula *trab*. For each *k*, the associated inhibition of I_Kr_ and I_CaL_ was computed using Eq. 1 and the IC50 data reported in Table 2. The minimisation of the cost function was performed using the scipy Python package. Based on experimental observations, the following constraints were put on the cubic surface during its fitting:

- ΔAPD_90_= 0 ms at baseline, i.e., *θ*_6_ = 0 ;
- ΔAPD_90_≤ −50 ms for 100% I_CaL_ block ;
- ΔAPD_90_≥ +320 ms for 100% I_Kr_ block ;
- 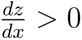, translating that an increase in I_Kr_ inhibition prolongs the APD_90_;
- 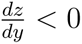, translating that an increase in I_CaL_ inhibition shortens the APD_90_.

When the constraints were not satisfied, the cost (Eq. 5) was multiplied by 100.

The goodness of fit of the cubic surface to the experimental points is plotted in Figure 10.

**Figure 10:**
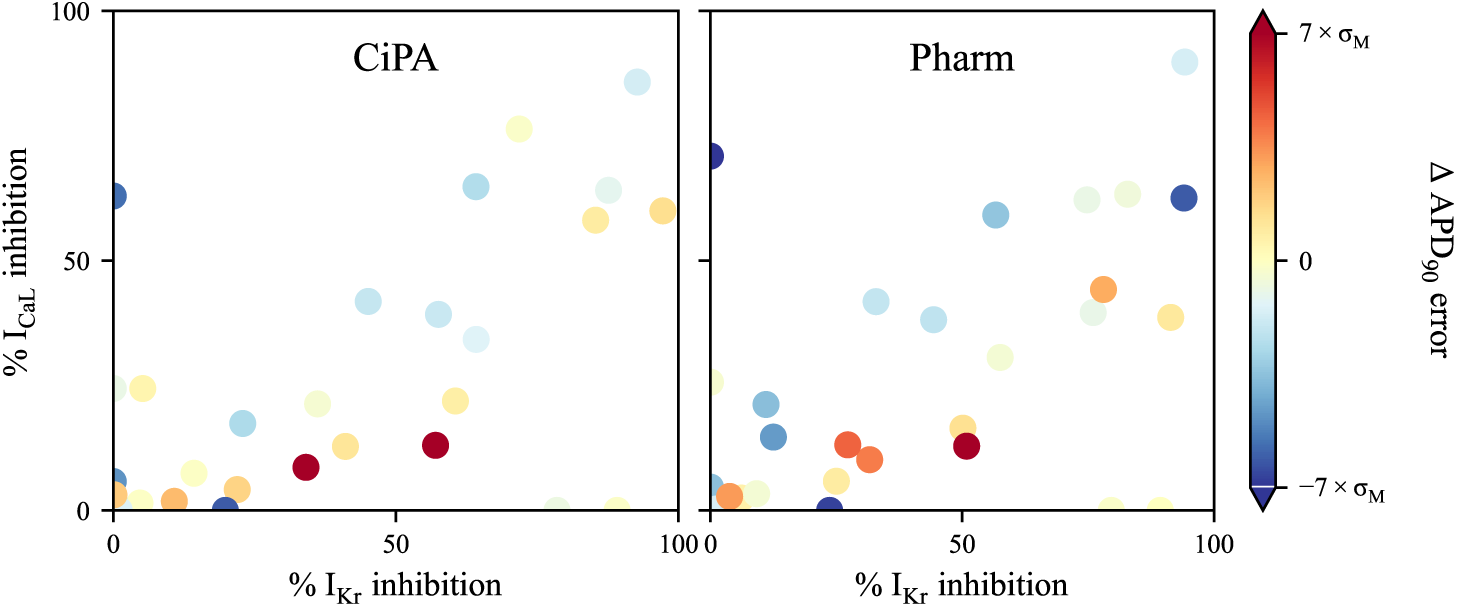
Error (Eq. 2) in the cubic surface compared with the experimental ΔAPD_90_ data, obtained using the CiPA (**left**) and the Pharm dataset (**right**).

## C Comparison of 2D maps using nominal concentrations and using measured drug concentrations in the bath solution (when available)

As reported in Table 1, drug concentrations in the bath solution were measured for some compounds, while they were not measured for other compounds. Note that for Clozapine, the drug concentration was measured in 7 trabeculae exposed to 0.3–3 *µ*M but not in the 4 trabeculae exposed to 0.3–30 *µ*M. In this section we compare whether fitting the cubic surface to nominal or measured drug concentrations makes any visual change to the resulting surface, and if simulating the drug-induced ΔAPD_90_ with nominal concentrations impacts our results.

The fitting of the cubic surface was repeated three times, using differently the data for drug concentration to locate the experimental data points on the 2-D map (Eq. 4):

- using the measured drug concentration in the bath solution for each trabecula under each tested drug condition, when the drug concentration was measured. Otherwise, the nominal concentration was used. In this case, ΔAPD_90_ is reported separately for each trabecula for each drug condition ;
- averaging the measured drug concentration for all trabeculae tested with the same nominal drug concentration, when the drug concentration was measured. Otherwise, the nominal drug concentration was used. The drug-induced ΔAPD_90_ effect was averaged over the trabeculae tested with the same nominal concentration ;
- using only the nominal drug concentrations. The drug-induced ΔAPD_90_ effect was averaged over the trabeculae tested with the same nominal concentration.

The experimental data points were placed on the 2-D map following the three methods, and the corresponding cubic surfaces were fitted through these data points. The fitting of the cubic surface was repeated with the CiPA and Pharm protocols, and the results are visualised in Figure 11.

The exact location of points was changed due to the measured concentrations in the bath solution not matching exactly with the nominal concentrations. Yet, the cubic surfaces fitted to the experimental points with the three methods described above were very similar. Therefore, the qualitative comparison of model predictions with the experimental data yields the same results when the nominal drug concentrations are used to compute the drug-induced inhibition of I_Kr_ and I_CaL_.

**Figure 11:**
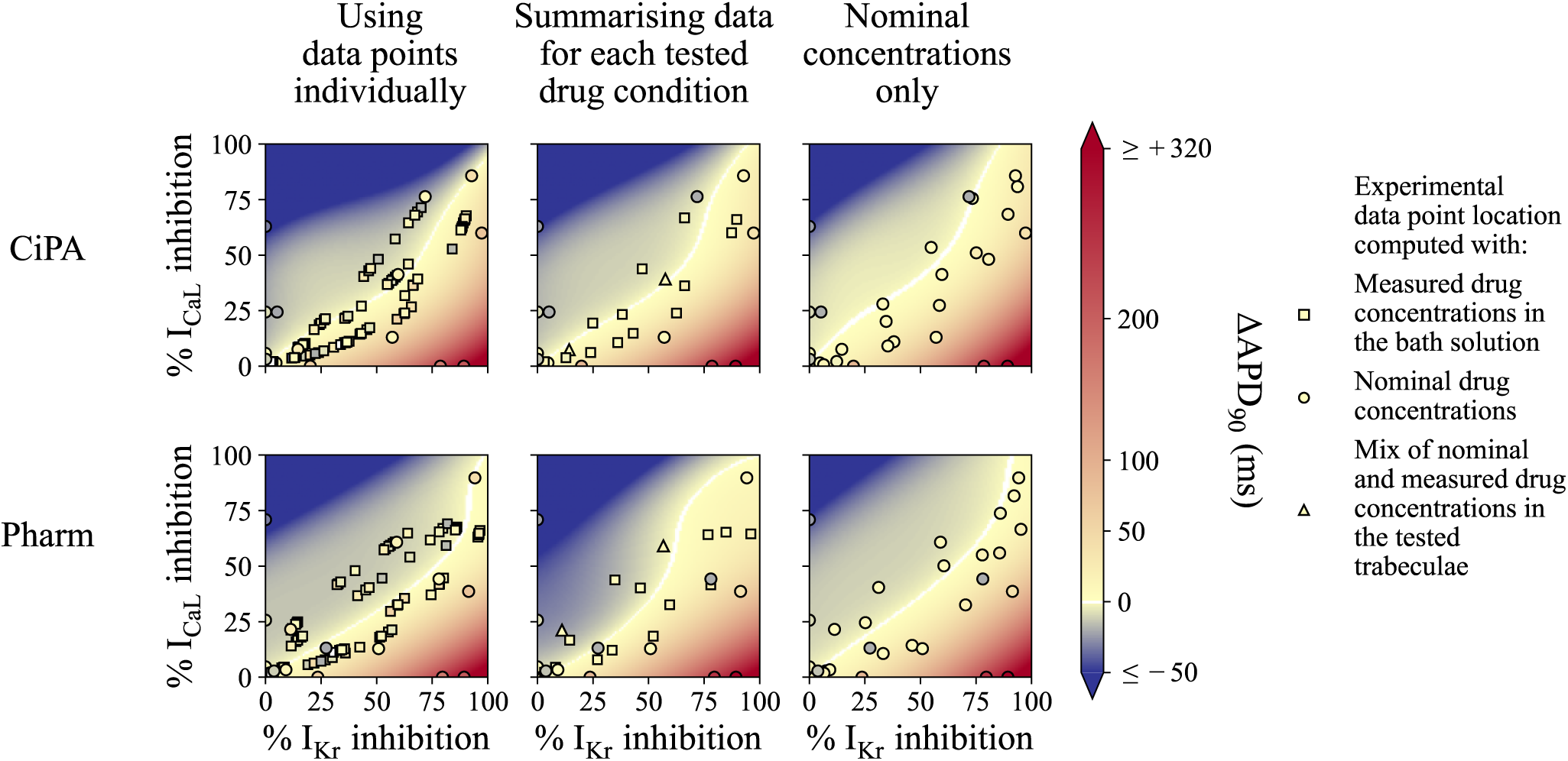
Impact of reporting drug concentrations as measured in the bath solution or nominal on the cubic surface and on the interpretation of the results in this study. Experimental ΔAPD_90_ measured ex-vivo under various drug conditions in human ventricular trabeculae, as a function of I_Kr_ and I_CaL_ inhibition and cubic surface approximating the experimental data points in the background. Each data point was placed with the current inhibition computed with Eq. 1 from drug concentrations and the drug IC50 for I_Kr_ and I_CaL_ (Table 2). **Left:** ΔAPD_90_ is reported with one point per tested drug condition per trabecula. The cubic surface was fitted to all the data points for all the trabeculae. **Middle:** A single point is plotted per tested drug condition. The data for drug concentration and drug-induced ΔAPD_90_ was averaged the trabeculae tested with the same nominal concentration. **Right:** A single point is ploted per tested drug condition, using the nominal drug concentration to compute the drug-induced current inhibition and subsequent location on the map. ΔAPD_90_ was averaged similarly to the middle panel. When measured drug concentrations were available only for some trabeculae tested with the same nominal drug concentration, the data point was plotted as a **triangle**.

To further support that the interpretation of our results were not sensitive to the discrepancy between measured drug concentrations in the bath solution and nominal drug concentrations, the scores were recomputed with all nominal concentrations, similarly to Figure 7. The results are similar to the results obtained with measured drug concentrations in the bath solution (when available), and the interpretation of our results therefore did not depend on the discrepancy between the measured and nominal drug concentrations.

## D Comparison of drug-induced ΔAPD_90_ with relative APD_90_ change from baseline as a percentage

To investigate whether drug-induced changes in APD_90_ should be reported in absolute or relative values, the correlation between baseline APD_90_ and response to drug perturbation was observed. For 15 trabeculae exposed to 1 *µ*M Verapamil, the absolute change in APD_90_ (ΔAPD_90_) was computed. From there, the relative change in APD_90_ was computed with the following equation:

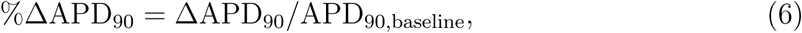

APD_90, baseline_ referring to the baseline APD_90_.

The scatter plot of ΔAPD_90_ and %ΔAPD_90_ against baseline APD_90_ is plotted in Figure 13. The ΔAPD_90_ and %ΔAPD_90_ showed not correlated with baseline APD_90_. Consistent results were obtained for all drugs and all drug concentrations.

Therefore, normalising ΔAPD_90_ would not improve the understanding of drug effect. Furthermore, in the clinic, changes in QT are measured in absolute, average prolongation of the QT interval by more than +5 ms being the limit of tolerance (ICH, 2006). At the cellular level, ΔAPD_90_ is more directly linked with the safety marker than %ΔAPD_90_. As a conclusion, the ΔAPD_90_ was used for the rest of this study.

To investigate further the importance of using ΔAPD_90_ over %ΔAPD_90_ in this study the results of the main text were repeated using the relative APD_90_ change expressed as a percentage of the baseline APD_90_ (%ΔAPD_90_), instead of ΔAPD_90_. The relative APD_90_ change was computed in each trabecula as:

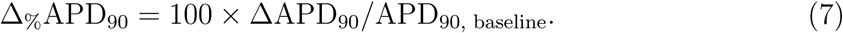

The experimental %ΔAPD_90_is plotted in Figure 14. The cubic surface was very similar to the cubic surface observed in Figure 4, granted the scalings differed between the two figures.

Predictions of %ΔAPD_90_ were also computed for the 11 AP models, and plotted in Figure 15. Very similar trends were observed in 2-D maps for ΔAPD_90_ and %ΔAPD_90_. Therefore, the choice of using ΔAPD_90_ over %ΔAPD_90_ did not impact the interpretation of the results presented in this study.

## E The TdP risk metric Q_net_ as a function of I_Kr_ and I_CaL_ inhibition

The CiPA initiative was established with the objective of developing an *in silico* model classifying drugs into three TdP risk categories (low, intermediate, high risk), providing a more specific safety assessment than the I_Kr_-centric guideline (Sager *et al*., 2014). One popular candidate, Q_net_, relies on the net charge flux over the repolarisation phase of one AP computed with the ORd-CiPA model (Li *et al*., 2019). Q_net_ is defined as the integral of the net currents that are active in the repolarisation phase over one AP at 0.5 Hz pacing after 1000 pre-paces defined as:

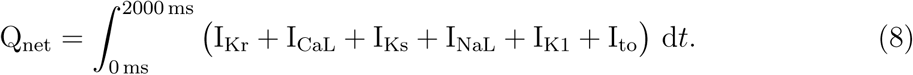

In this section, we compare Q_net_ as a function of I_Kr_ and I_CaL_ inhibition with experimental ΔAPD_90_ measurements.

As with the AP models, Q_net_ was computed with the ORd-CiPA model for 101 × 101 = 10, 201 combinations of I_Kr_ and I_CaL_ inhibition. The reduction of I_Kr_ was modelled by applying a multiplying factor to the maximal conductance of I_Kr_, as in Section 2.4.1, although in the original methods of Li *et al*., the I_Kr_ inhibition by drugs is modelled with the dynamic hERG binding model (Li *et al*., 2017). Note that Li *et al*. classified compounds into the TdP risk categories based on the average Q_net_ computed at 1–4 times their maximal effective free therapeutic concentration. Nevertheless, a qualitative interpretation of the 2-D map remains possible.

Pixels of the 2-D map were colored based on the TdP risk category corresponding to Q_net_ obtained with the ORd-CiPA model after inhibition of the ionic currents. As in (Li *et al*., 2019), Q_net_ values greater than 0.0671 *µ*C.*µ*F^−1^ were classified as low risk (green), Q_net_ between 0.0581 *µ*C.*µ*F^−1^ and 0.0671 *µ*C.*µ*F^−1^ as intermediate risk (blue), and Q_net_ lower than 0.0581 *µ*C.*µ*F^−1^ as high risk.

The resulting 2-D map is shown in Figure 16.

Interestingly, the decrease in Q_net_ (increase in TdP risk) induced by I_Kr_ inhibition is mitigated by I_CaL_ inhibition, with a higher sensitivity to I_CaL_ inhibition than ΔAPD_90_ predicted by the ORd-CiPA model. For example, 50% I_Kr_ inhibition and 75% I_CaL_ inhibition yields a Q_net_ value classified into the low TdP risk category, while the predicted ΔAPD_90_ is +73 ms.

The shape of the 2-D map of Q_net_ is similar to that of the 2-D map of ΔAPD_90_ predicted by the TP-M model (Figure 5). Furthermore, qualitatively similar mitigation of I_Kr_ inhibition by I_CaL_ inhibition was observed between Q_net_ predictions and ΔAPD_90_ observed experimentally (Figure 4).

**Table 4:** A summary of trabeculae recordings for average APD_90_ at baseline and drug-induced APD_90_ change from baseline (ΔAPD_90_). SEM: Standard error of the mean. SD: Standard deviation.

| Drug | Mean baseline<br>APD<sub>90</sub> (SD) in<br>ms | Nominal<br>drug conc<br>(μM) | Mean<br>ΔAPD<sub>90</sub><br>(SEM), in ms |
| --- | --- | --- | --- |
| Chlorpromazine | 299 (36) | 0.3 | +9 (10) |
|  |  | 1 | +18 (8) |
|  |  | 3 | +24 (11) |
| Clozapine | 324 (51) | 0.3 | +8 (5) |
|  |  | 1 | +10 (7) |
|  |  | 3 | +10 (7) |
|  |  | 30 | +15 (9) |
| Dofetilide | 317 (51) | 0.001 | +20 (5) |
|  |  | 0.01 | +82 (8) |
|  |  | 0.1 | +256 (21) |
|  |  | 0.2 | +318 (33) |
| Fluoxetine | 271 (36) | 0.3 | +10 (4) |
|  |  | 1 | +6 (7) |
|  |  | 3 | -2 (5) |
| Mesoridazine | 334 (60) | 0.04 | -2 (7) |
|  |  | 0.25 | +2 (4) |
|  |  | 10 | +21 (2) |
| Nifedipine | 336 (55) | 0.003 | +7 (4) |
|  |  | 0.03 | -5 (6) |
|  |  | 0.3 | -24 (6) |
| Quinidine | 302 (53) | 0.1 | +6 (5) |
|  |  | 1 | +8 (5) |
|  |  | 10 | +37 (7) |
| Thioridazine | 307 (48) | 0.012 | +15 (9) |
|  |  | 0.6 | +16 (20) |
|  |  | 2 | +6 (14) |
| Verapamil | 349 (61) | 0.01 | -15 (4) |
|  |  | 0.1 | -19 (5) |
|  |  | 1 | -20 (10) |

**Table 5:** Data summary of IC_50_ values for hERG and Ca_V_1.2 block potency of the Clozapine.

| Drug | hERG<br>block IC50<br>(μM)/h | Ca<sub>v</sub>1.2<br>block IC50<br>(μM)/h | Source |
| --- | --- | --- | --- |
| Clozapine | 1.978 / 0.94 | 4.378 / 0.93 | CiPA prot. |
|  | 2.123 / 1.05 | 1.676 / 0.75 | Pharm prot. |
|  | 2.300 / 0.97 | 3.600 / 1 | (Kramer *et al.*, 2013) |
|  |  | 5.490 / 0.94 | (Li *et al.*, 2019) |
|  | 2.310 / 1 | 3.555 / 1 | One Million Solutions in Health |
|  | 1.639 / 1 | 3.600 / 1 | (Llopis-Lorente *et al.*, 2020) |

**Table 6:** Data summary of IC_50_ values for hERG and Ca_V_1.2 block potency of Dofetilide.

| Drug | hERG<br>block IC50<br>(μM)/h | Ca<sub>v</sub>1.2<br>block IC50<br>(μM)/h | Source |
| --- | --- | --- | --- |
| Dofetilide | 0.033 / 1.17 |  | CiPA prot. |
|  | 0.029 / 1.10 | 332 / 1 | Pharm prot. |
|  | 0.030 / 1.2 | 26.7 / 1 | (Kramer *et al.*, 2013) |
|  | 0.0061 / 1.08 | 44.5 / 3.6 | CiPA GitHub |
|  | 0.005 / 1 | 60 / 1 | (Mirams *et al.*, 2011) |
|  | 0.002 / 1 |  | (Redfern *et al.*, 2003) |
|  | 0.002 / 1 |  | (Crumb Jr *et al.*, 2016) |
|  | 0.037 / 2.1 | 182 / 1 | (Okada *et al.*, 2015) |
|  | 0.025 / 1 | 26.7 / 1 | (Llopis-Lorente *et al.*, 2020) |

**Table 7:**
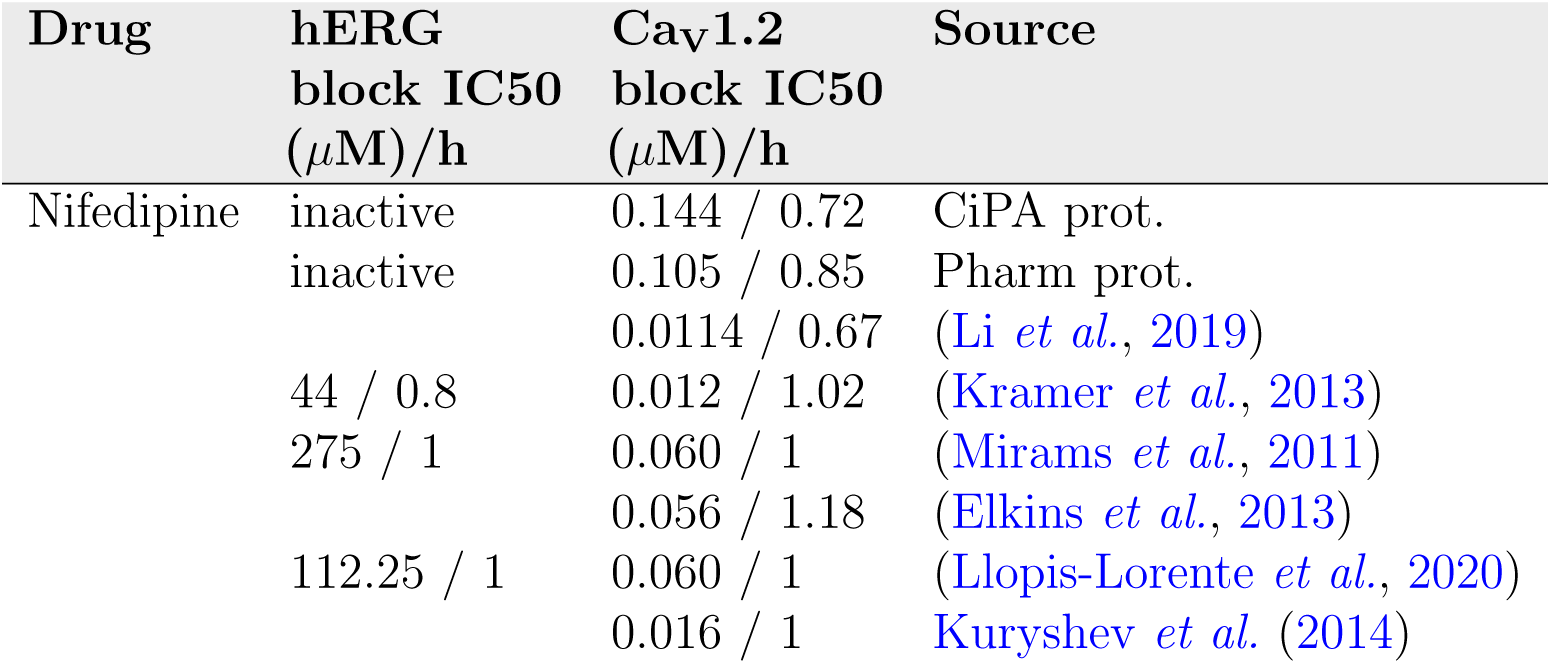
Data summary of IC_50_ values for hERG and Ca_V_1.2 block potency of Nifedipine.

**Table 8:** Data summary of IC_50_ values for hERG and Ca_V_1.2 block potency of Quinidine.

| Drug | hERG<br>block IC<sub>50</sub><br>(μM)/h | Ca<sub>v</sub>1.2<br>block IC<sub>50</sub><br>(μM)/h | Source |
| --- | --- | --- | --- |
| Quinidine | 0.820 / 1.43 | 6.680 / 1 | CiPA prot. |
|  | 0.966 / 1.01 | 20.849 / 0.63 | Pharm prot. |
|  | 2.371 / 1.71 |  | (Elkins *et al.*, 2013) |
|  | 0.3 / 1 | 15.6 / 1 | (Mirams *et al.*, 2011) |
|  | 0.72 / 1.06 | 6.4 / 0.68 | (Kramer *et al.*, 2013) |
|  | 0.3 / 1 |  | (Crumb Jr *et al.*, 2016) |
|  | 0.658 / 1 | 8.1 / 0.8 | (Okada *et al.*, 2015) |
|  | 0.986 / 0.84 | 51.59 / 0.59 | CiPA GitHub |
|  | 0.890 / 1 | 8.866 / 0.912 | (Llopis-Lorente *et al.*, 2020) |

**Table 9:** Data summary of IC_50_ values for hERG and Ca_V_1.2 block potency of Verapamil.

| Drug | hERG<br>block IC<sub>50</sub><br>(μM)/h | Ca<sub>v</sub>1.2<br>block IC<sub>50</sub><br>(μM)/h | Source |
| --- | --- | --- | --- |
| Verapamil | 0.570 / 1.67 | 0.310 / 1 | CiPA prot. |
|  | 0.273 / 0.98 | 1.381 / 0.72 | Pharm prot. |
|  | 0.296 / 0.94 | 0.202 / 1.10 | CiPA GitHub |
|  | 0.250 / 0.89 | 0.200 / 0.8 | (Kramer *et al.*, 2013) |
|  | 0.677 / 1.43 | 2.685 / 0.61 | (Elkins *et al.*, 2013) |
|  | 0.143 / 1 | 0.100 / 1 | (Mirams *et al.*, 2011) |
|  | 0.7 / 1 | 0.1 / 1 | (Crumb Jr *et al.*, 2016) |
|  | 0.212 / 1 | 0.347 / 1.08 | (Okada *et al.*, 2015) |
|  | 0.499 / 1.1 | 0.201 / 1.1 | (Llopis-Lorente *et al.*, 2020) |

**Figure 12:**
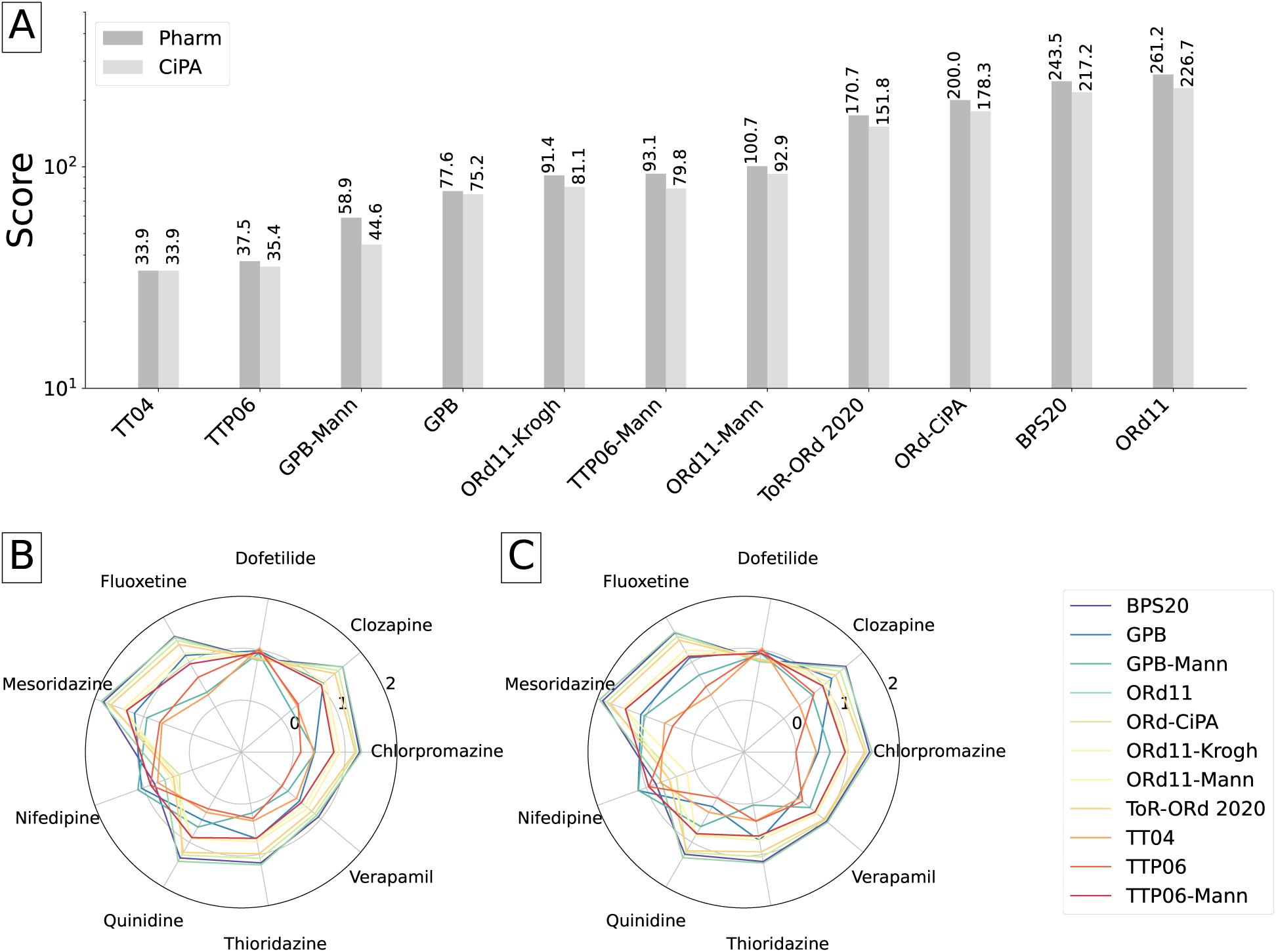
Comparison of the abilities of human ventricular AP models to reproduce the APD_90_ response to I_Kr_ and I_CaL_ inhibition observed *ex vivo*, using only nominal drug concentrations as inputs for simulations. The error metric was computed from Eq. 2. **A:** The error measure was summed over all the drugs used in this study, when using the Pharm and CiPA protocols to compute the reduction of ionic currents by drugs. For each model, two bar plots were plotted, to compare the predictive power of models with the Pharm (left bar) and the CiPA (right bar) datasets. **B and C:** Detail of the error measures associated to each of the drugs with the CiPA and Pharm datasets, respectively, for each model. The log_10_ of the error measure is plotted along the radial-axis.

**Figure 13:**
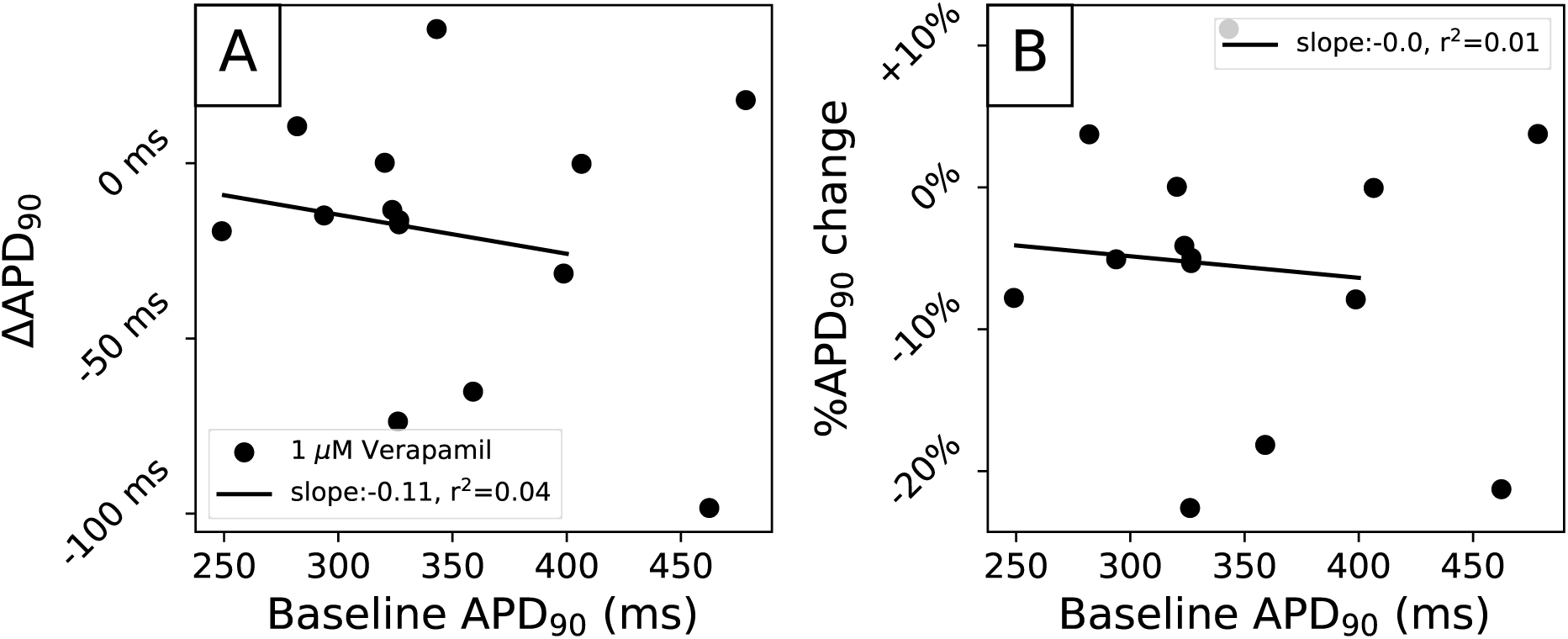
Correlation between baseline APD_90_ and Verapamil-induced perturbations, measured as absolute change in APD_90_ (**A**) and as relative change in APD_90_ (**B**). No correlation was found.

**Figure 14:**
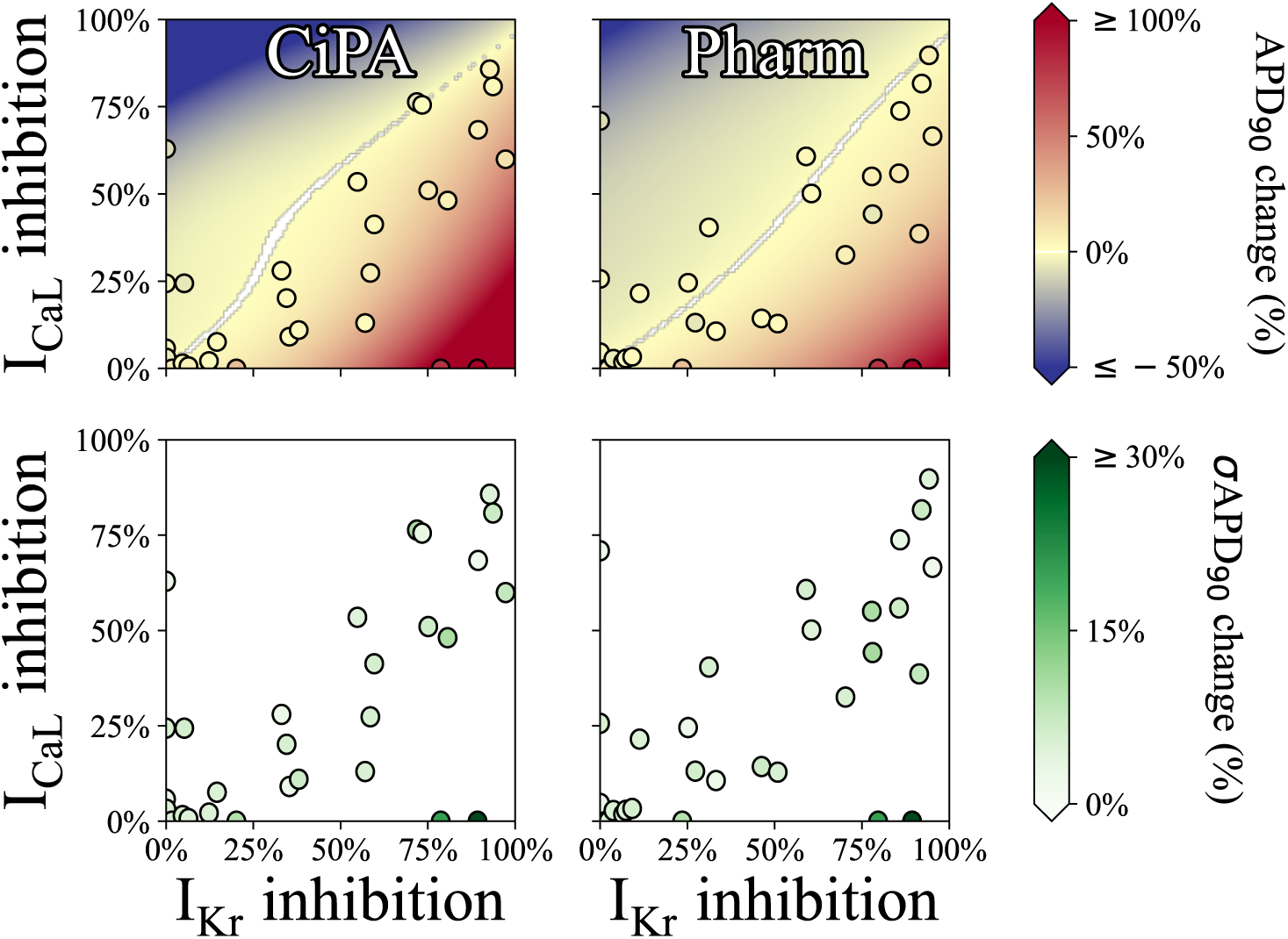
Experimental %ΔAPD_90_ measured ex-vivo under various drug conditions in human ventricular trabeculae, as a function of I_Kr_ and I_CaL_ inhibition and cubic surface approximating the experimental data points in the background. I_Kr_ and I_CaL_ inhibition were computed using the Hill equation (Eq. 1), with the CiPA (left) and Pharm (right) datasets (Table 2) and nominal drug concentrations (Table 1).

**Figure 15:**
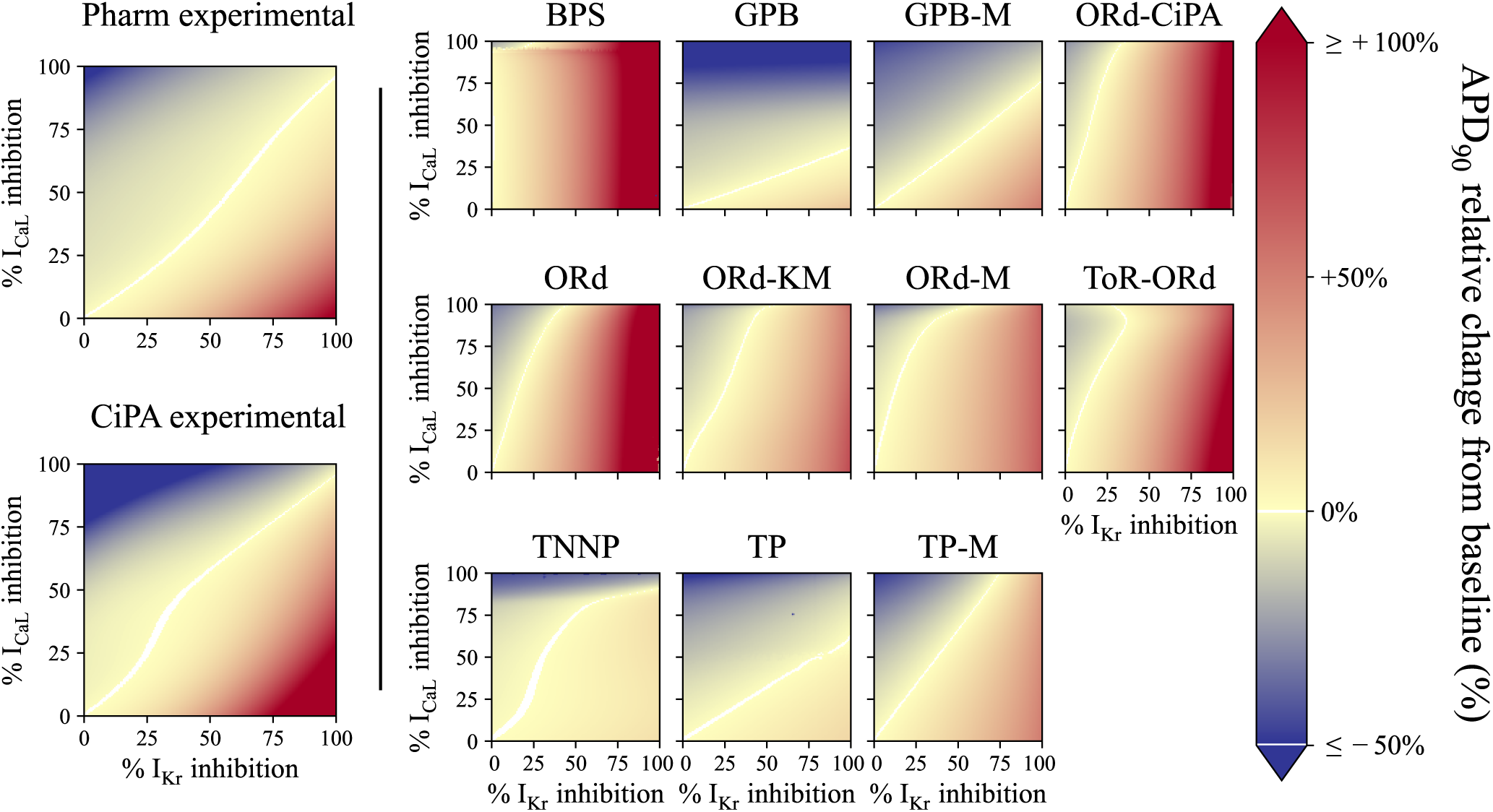
2-D maps of simulated %ΔAPD_90_after I_CaL_ and I_Kr_ inhibition. The colour scale indicates shortening of APD_90_ (i.e., Δ_%_APD_90_ *<* 0 ms) for colours towards dark blue, and APD_90_ prolongation (i.e., Δ_%_APD_90_ *<* 0 ms) for colours towards red. ΔAPD_90_ values below −50 ms and above +200 ms were set to dark blue and red, respectively, for better visualisation. For I_Kr_ and I_CaL_ inhibition leading to −1 ms *<* ΔAPD_90_ *<* +1 ms, the pixel is coloured in white.

**Figure 16:**
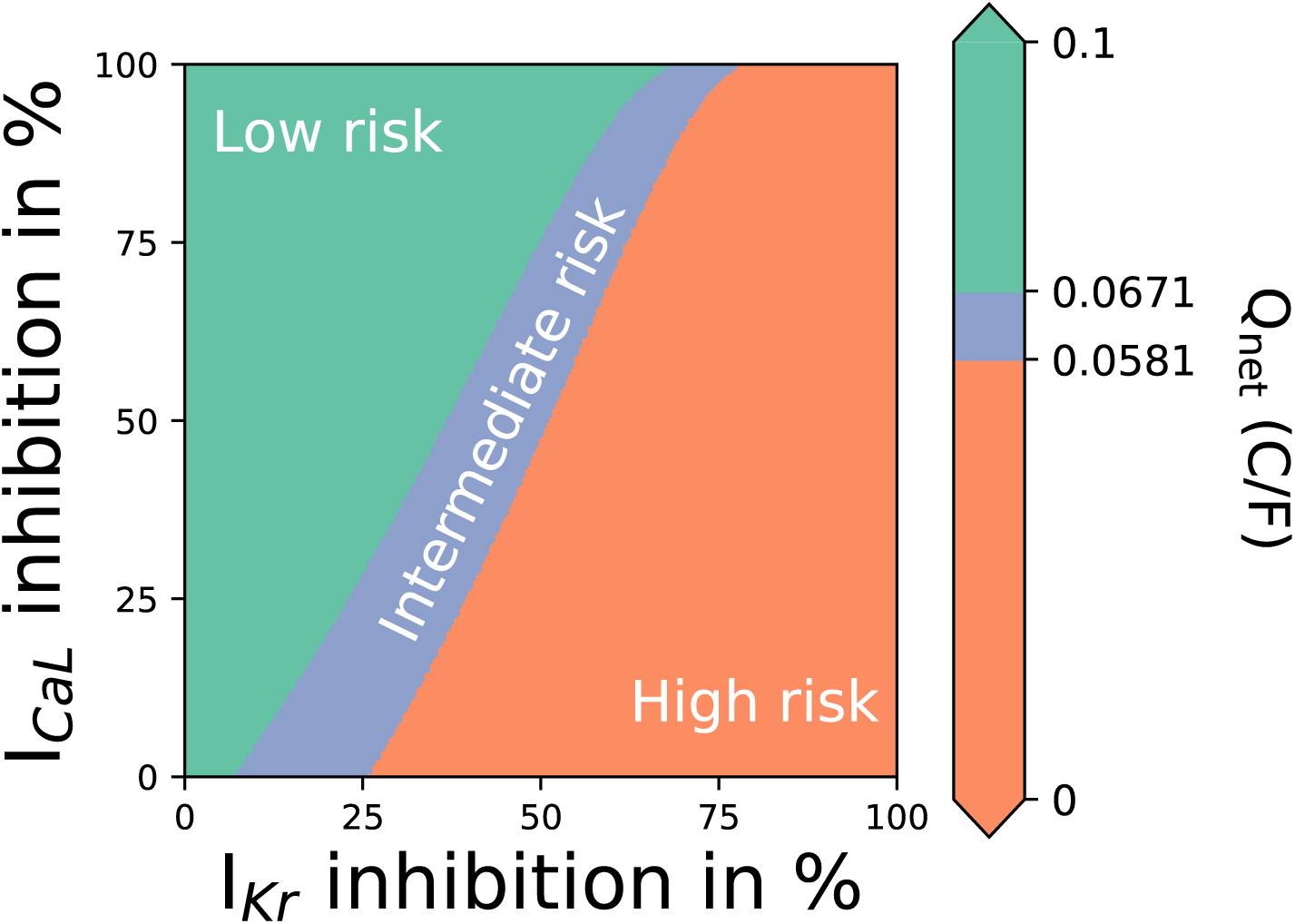
Q_net_ computed with the ORd-CiPA model, for various combinations of I_Kr_ and/or I_CaL_ inhibition. The Torsade risk thresholds are defined in Li *et al*. (2019). Note that I_Kr_ inhibition was computed as plain reduction of the maximal conductance of I_Kr_ (Section 2.4.1) instead of the dynamic hERG binding model used in the original study (Li *et al*., 2019).

